# Sleep promotes downward firing rate homeostasis

**DOI:** 10.1101/827832

**Authors:** Alejandro Torrado Pacheco, Juliet Bottorff, Gina G. Turrigiano

## Abstract

Homeostatic plasticity is hypothesized to regulate neuronal activity around a stable set point to compensate for learning-related plasticity. This regulation is predicted to be bidirectional but only upward firing rate homeostasis (FRH) has been demonstrated *in vivo*. We combined chronic electrophysiology in freely behaving animals with a protocol that induces robust plasticity in primary visual cortex (V1) to induce downward FRH and show that neurons bidirectionally regulate firing rates around an individual set point. Downward FRH did not require N-methyl-D-aspartate receptor (NMDAR) signaling and was associated with homeostatic scaling down of synaptic strengths. Like upward FRH, downward FRH was gated by vigilance state, but in the opposite direction: it occurred during sleep and not during wake. In contrast, FR changes associated with Hebbian plasticity happened independently of sleep and wake. Thus, we find that sleep’s impact on neuronal plasticity depends on the particular forms of plasticity that are engaged.

## INTRODUCTION

Proper functioning of neocortical networks requires that they be simultaneously plastic and stable, and these demands necessitate the expression of a variety of Hebbian and homeostatic mechanisms that modify synaptic connections and neuronal firing rates (FRs) (Abbott and Nelson, 2000; Turrigiano and Nelson, 2004). Hebbian mechanisms that strengthen or weaken synaptic strengths as a function of correlated spiking are widely considered to provide the basis for long-term storage of information in the brain, but are also theorized to be intrinsically destabilizing when left unchecked. Modeling studies support this idea and generally come to the conclusion that neuronal networks require compensatory forces that stabilize activity during experience-dependent plasticity and learning (Miller and MacKay, 1994; Tetzlaff et al., 2011; Litwin-Kumar and Doiron, 2014). Homeostatic mechanisms are hypothesized to provide this balance by regulating neuronal activity around a stable FR set point, a process known as firing rate homeostasis (FRH; Turrigiano and Nelson, 2004). Upward firing rate homeostasis has now been convincingly demonstrated in the mammalian visual system *in vivo* using a variety of approaches (Kaneko et al., 2008; Keck et al., 2013; Hengen et al., 2013; Barnes et al., 2015; Hengen et al., 2016), and strikingly is gated by behavioral state so that it occurs exclusively during active wake (Hengen et al., 2016). Here we ask whether firing rate homeostasis is bidirectional as theory predicts, and whether upward and downward homeostatic plasticity are gated similarly by sleep/wake states.

That sleep and wake play a key role in regulating brain plasticity is widely accepted, but experiments designed to understand the details of this regulation have yielded contradictory results (Puentes-Mestril and Aton, 2017). A prominent hypothesis (the synaptic homeostasis hypothesis, SHY) posits that a net potentiation of synaptic strengths and FRs occurs during waking, while the function of sleep, and in particular of non-rapid-eye-movement (NREM) sleep, is to enable homeostatic down-regulation of these parameters (Tononi and Cirelli, 2003; 2014). Evidence in support of SHY has accumulated, but many results inconsistent with it have also been reported, and it remains a controversial theory (Frank and Cantera, 2014). For instance, neuronal FRs have been found to be higher after wake in the frontal cortex (Vyazovskiy et al., 2009) but not different across sleep or wake in visual cortex (Hengen et al., 2016), while other experiments find that sleep- and wake-driven changes in FR are not uniform across neuronal populations (Watson et al., 2016). Importantly, these findings focused on FR dynamics in the absence of plasticity induction. Here we combine our ability to track the activity of individual neurons with a protocol that triggers robust plasticity to specifically dissect the impact of sleep and wake states on downward FRH.

To induce and observe downward FRH, we performed chronic electrophysiology in monocular visual cortex (V1m) of freely behaving rats undergoing monocular deprivation (MD) followed by eye re-opening (ER) on day 5 of MD, a protocol known to increase activity within V1 (Toyoizumi et al., 2014). We found a robust increase in FR within 24 hours of ER, followed by a gradual return of the firing rates of individual neurons to their pre-MD baseline levels. Using blockade of N-methyl-D-aspartate receptors (NMDARs) and measurements of synaptic strengths in acute slices we show that the FR overshoot following ER is consistent with Hebbian plasticity mechanisms, while the subsequent recovery of activity is consistent with homeostatic synaptic scaling. Interestingly, this downward FRH only occurred during sleep-dense epochs, or during periods of extended sleep, in contrast to upward FRH which happens during wake (Hengen et al., 2016). The rules governing the expression and interaction of Hebbian and homeostatic mechanisms *in vivo* remain unknown (Keck et al., 2017), so we wondered whether Hebbian plasticity was also gated by sleep and wake. When we analyzed the state dependence of the MD-induced reduction in FR, which is mainly driven by Hebbian long-term depression (LTD), we found that it unfolded independently of sleep and wake states. Our data support a model in which downward Hebbian plasticity happens independently of sleep and wake, while homeostatic mechanisms are differentially gated by sleep and wake states depending on the direction of compensation.

## RESULTS

Bidirectional homeostatic regulation of FRs has not been demonstrated *in vivo*, and the role of sleep in its induction is unknown. Further, it is unclear how the homeostatic and Hebbian mechanisms that interact to modify neuronal activity are integrated *in vivo*, and whether sleep and wake states play a role in their orchestration. To examine these questions we recorded single-unit activity from V1m of freely behaving rats undergoing MD and then ER, and analyzed the behavior of individual regular spiking (putative pyramidal) neurons that could be continuously recorded during this paradigm. We monitored sleep and wake states throughout these multi-day recordings to assess their impact on plasticity, and paired this with pharmacology and *ex vivo* synaptic interrogation to tease apart the impact of sleep and wake states on different plasticity mechanisms.

### Eye re-opening after MD causes an overshoot in FRs followed by homeostatic recovery

Prolonged MD in rats induces bi-phasic changes in activity in V1 *in vivo* (Hengen et al., 2016). To determine whether the homeostatic regulation of FRs is bi-directional, we sought to cause an overshoot in FR above baseline levels in neurons in V1m. To achieve this, we bilaterally recorded single-unit activity in V1m continuously for up to 11 days in freely behaving young rats (P24-35) undergoing MD after 3 days of baseline and eye re-opening (ER) on the 5^th^ day of MD, when the FRs of V1m neurons have on average returned to baseline levels. In this experimental design neurons recorded from the hemisphere ipsilateral to the manipulated eye constitute a within-animal control (control hemisphere). We reproduced our previous finding that FRs drop after 2 days of MD and subsequently recover to baseline. ER on day 5 of MD caused a similar bi-phasic pattern of change in the opposite direction, namely an overshoot of activity above baseline levels, followed by a downward recovery (Figure 1A-D; Figure S1). To analyze this further we focused on the time around ER, taking the 12 hours prior to ER (day 4 of MD, MD4) as our new baseline (Figure S1). ER increased firing in individual neurons with very different baseline firing rates, followed by a slow recovery back to baseline (Figure 1B); In contrast neurons in the control hemisphere were unaffected by ER (Figure 1A). The same pattern was observed when firing rates across the population were normalized and averaged (Figure 1C, D).

**Figure 1:**
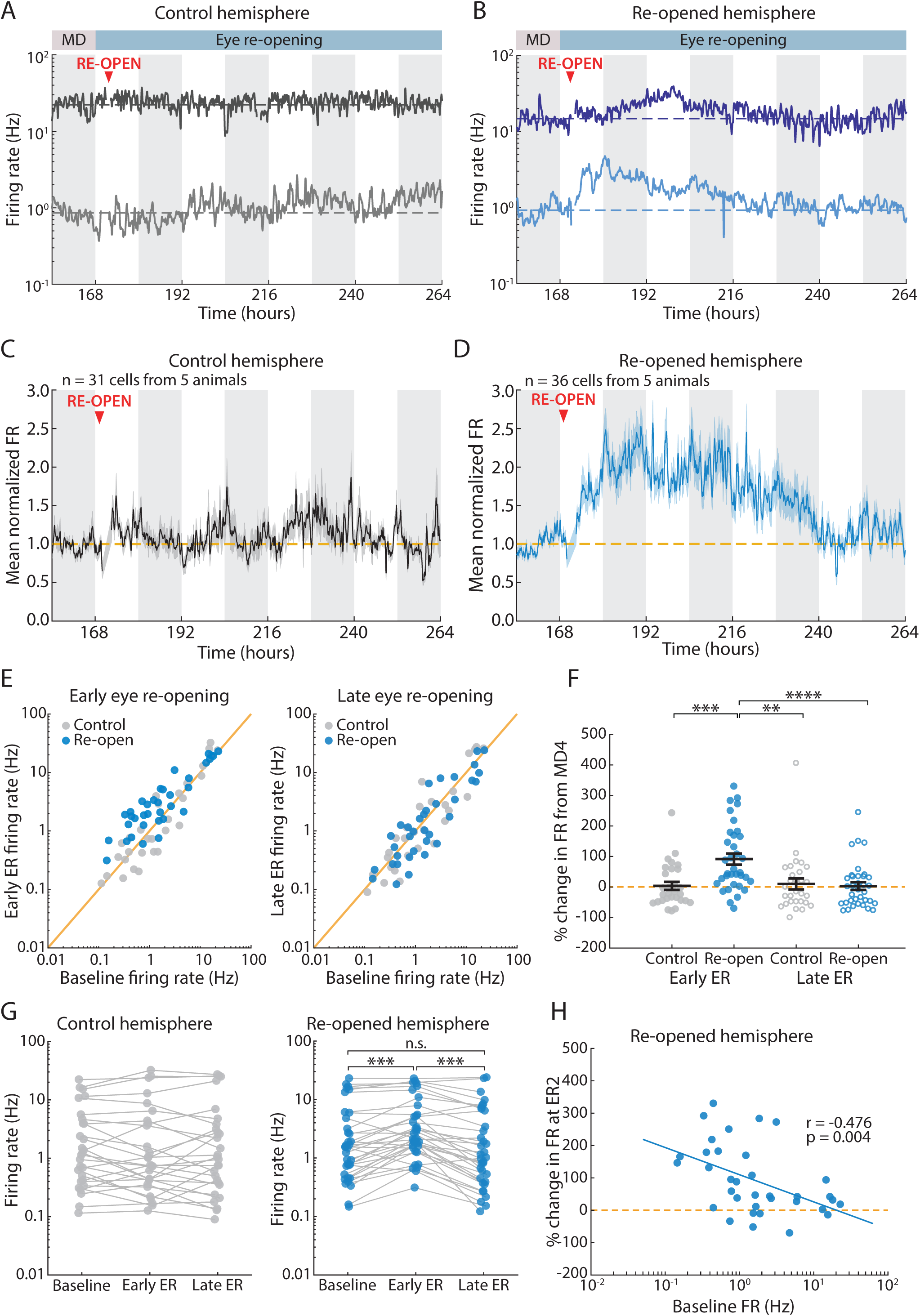
Eye re-opening after MD causes an overshoot in FRs followed by homeostatic recovery. **A**. Example neurons recorded continuously from the control hemisphere for 5 days. Dashed line indicates the mean FR of the neuron during the baseline (MD4) period. **B**. Example neurons recorded continuously from the re-opened hemisphere for 5 days. Here eye re-opening causes an increase in FR followed by a homeostatic recovery. Dashed line indicates baseline FR. Artifact due to unplugging animals for ER surgery has been removed. **C, D**. Average FR traces for all neurons recorded in control and re-opened hemispheres, normalized to baseline (MD4) for each cell. Dashed line indicates baseline FR. Artifact due to unplugging animals for ER surgery has been removed. **E**. Comparison of individual neuronal FRs between baseline (MD4) and early ER (left) or late ER (right). Each dot is the mean FR for a given neuron averaged over the corresponding 12-hour period. Yellow line indicates unity (no change in FR). **F**. Percent change in FR from baseline at early and late ER for each control and re-opened hemisphere neuron. Black lines indicate mean ± SEM. Control n = 29; re-open n = 35; Kruskal-Wallis test (p < 0.0001) with Tukey-Kramer post-hoc, ** p = 0.003, *** p = 0.002, **** p = 0.0004. **G**. Mean FR of every cell at baseline (MD4), early ER and late ER. Each dot represents mean FR for one cell over the corresponding 12-hour period, and mean FRs for the same cell are connected via solid lines. Control, n = 31; Re-open, n = 35; Wilcoxon sign-rank test with Bonferroni correction, *** p< 0.001. **H**. Correlation between mean baseline FR and percent change in FR at early ER. Solid line represents linear fit. Correlation estimated using Pearson’s method, n = 35, *r* = −0.476, p = 0.0039.

V1 RSUs have mean FRs that span several orders of magnitude. To compare the pattern of activity across the population, we compared the baseline FR (MD4) to FRs on day 2 or day 4 after ER (early ER, ER2; late ER, ER4) for each neuron (Fig. 1E-H). On ER2 most neuronal FRs from the re-opened hemisphere were elevated compared to controls (Early eye re-opening), while the two distributions were similar by ER4 (Late eye re-opening, Figure 1E). We quantified the changes at these time-points by computing the percent change in FR from baseline (Figure 1F). The FRs of re-opened hemisphere neurons were significantly higher on ER2 (114 ± 23% change from MD4), but by ER4 the mean change from MD4 was near 0% (2 ± 13%), and similar to the mean change for control neurons (10 ± 18%). Examining the behavior of individual neurons across time showed a significant change only in the re-opened hemisphere, where most neurons increase and then decrease firing again (Figure 1G). Notably, we used a bootstrap analysis to show that the mean change between baseline and ER4 of close to 0% in the re-opened hemisphere can best be explained if each neuron’s FR returns close to its initial baseline rate (Figure S2), suggesting (in agreement with upward FRH, Hengen et al., 2016) that neurons regulate firing around an individual setpoint. Finally we asked whether the magnitude of the FR potentiation induced by ER was dependent on the initial baseline FR. We found a significant negative correlation between baseline FR and percent change in FR at ER2 (*r* = −0.476, *p* = 0.004) in the re-opened hemisphere only, indicating that low FR neurons show stronger potentiation than high FR neurons (Figure 1H).

These data demonstrate that neuronal FRs are homeostatically regulated *in vivo* in a bidirectional manner, and that during downward firing rate homeostasis neurons return close to an individual FR set-point.

### ER-induced FR overshoot, but not downward recovery, is NMDAR-dependent

Long-lasting changes in FR can be driven by various plasticity mechanisms whose action requires different pathways and receptors. We set out to test whether the changes caused by ER were dependent on the activity of N-methyl-D-aspartate receptors (NMDARs), which are required for many forms of Hebbian plasticity (Malenka and Bear, 2004), but not for homeostatic synaptic scaling (Turrigiano et al., 1998). To this effect we used systemic injections of 3-(2-Carboxypiperazin-4-yl)propyl-1-phosphonic acid (CPP), a potent NMDAR antagonist that has been shown to block induction of LTP up to 24 hours after administration (Villarreal et al., 2002; Sato and Stryker, 2008; Toyoizumi et al., 2014). When CPP was injected at the time of ER, it completely blocked the increase in FR (Figure 2A, C). We found no change in the distribution of FRs between MD4 and ER2 in this case, and a slight decrease between ER2 and ER4 (Figure 2E). Neurons in this condition showed no percent change in FR at ER2 (Figure 2F; 6 ± 16%), and a slight decrease from ER2 to ER4 (Figure 2F; – 32 ± 6%).

**Figure 2:**
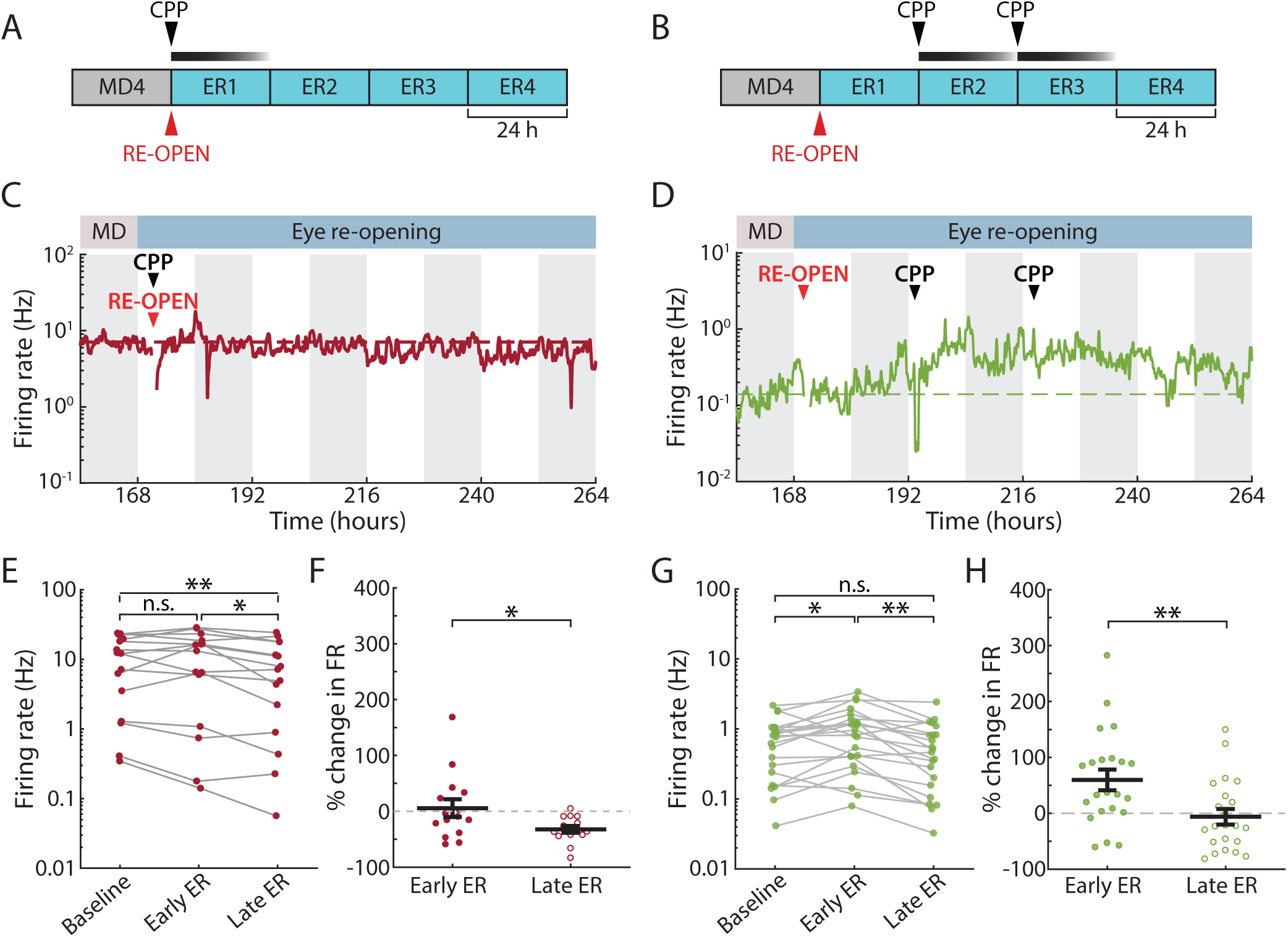
ER-induced FR overshoot, but not downward recovery, is NMDAR-dependent. **A, B**. Experiment schematic: CPP (15 mg/kg) was injected subcutaneously at the time of eye re-opening (A), or twice after that, at 24-hour intervals (B). **C, D**. Example FRs of neurons recorded in each of the CPP experiments. **E**. Change in FR from baseline to early ER to late ER for neurons recorded in the first CPP condition (injection at time of ER). Wilcoxon sign-rank test with Bonferroni correction, n = 15, ** p = 0.0026; * p = 0.0251. **F**. Percent change in FR from baseline in the first CPP condition. One-sample t-test comparing to mean = 0, ER2 p = 0.723, ER4 p < 10^−4^; two-sample t-test, n = 15, * p = 0.0293. **G**. As in E, but for the second CPP condition (two injections, 24 hr and 48 hr after ER). Wilcoxon sign-rank test with Bonferroni correction, n = 22, * p = 0.0269; ** p = 0.0014. **H**. As in F, but for second CPP condition. One-sample t-test comparing to mean = 0, ER2 p = 0.003, ER4 p = 0.657; two-sample t-test, n = 22, ** p = 0.0058.

Many homeostatic plasticity mechanisms do not require NMDAR activity (Turrigiano, 2008). If downward FRH is due to NMDAR-independent homeostatic plasticity, then if we first allowed ER to potentiate firing and then administered CPP we should see little effect on the downward homeostatic recovery of FRs. We tested this by injecting CPP twice, at 24-hour intervals, starting 24 hours after ER (Figure 2B). Despite a small and acute depressive effect of CPP on activity immediately following administration in both control and deprived hemispheres (Figure S3), we observed a normal ER-induced increase prior to CPP administration (FR change at ER2: 60 ± 18%), and then subsequent recovery of firing in this condition, similar to un-injected animals (Figure 2D, G, H).

We conclude that the increase in FR following ER is not simply due to increased sensory drive, but it is instead an active process that requires NMDAR-dependent plasticity. Conversely the subsequent downward firing rate homeostasis is independent of NMDAR activity, consistent with it being driven by homeostatic plasticity mechanisms.

### Downward FR recovery is associated with synaptic scaling down

Having established that the downward firing rate homeostasis is NMDAR-independent, we wished to know whether it is accompanied by synaptic scaling down, one of the principal forms of homeostatic plasticity within V1 (Turrigiano et al., 1998; Turrigiano, 2008). We performed MD and ER on rats as described above, and recorded miniature excitatory postsynaptic currents (mEPSCs) from layer 2/3 pyramidal neurons in V1m either 24 hours (ER2) or 72 hours (ER4) after ER (Figure 3A). Neurons in the control hemispheres of the same animals were used as controls. Example recordings and mean event amplitudes are shown in Figure 3B. While mEPSC amplitudes were stable in the control condition (ER2: 10.82 ± 0.35 pA; ER4: 10.32 ± 0.24 pA), neurons in the re-opened hemisphere showed a significant increase in mEPSC amplitudes at ER2 (11.56 ± 0.21 pA), and then a depression by ER4 (10.51 ± 0.23 pA; Figure 3C). There were no changes in mEPSC frequency, passive neuronal properties, or waveform kinetics for any conditions (Figure S4). Analysis of the cumulative distribution function (CDF) for all recorded mEPSC events revealed a significant shift to higher amplitudes in the ER2 re-opened condition, compared to both ER4 re-opened (*p* < 10^−5^) and ER4 control (*p* < 10^−6^; Figure 3D).

**Figure 3:**
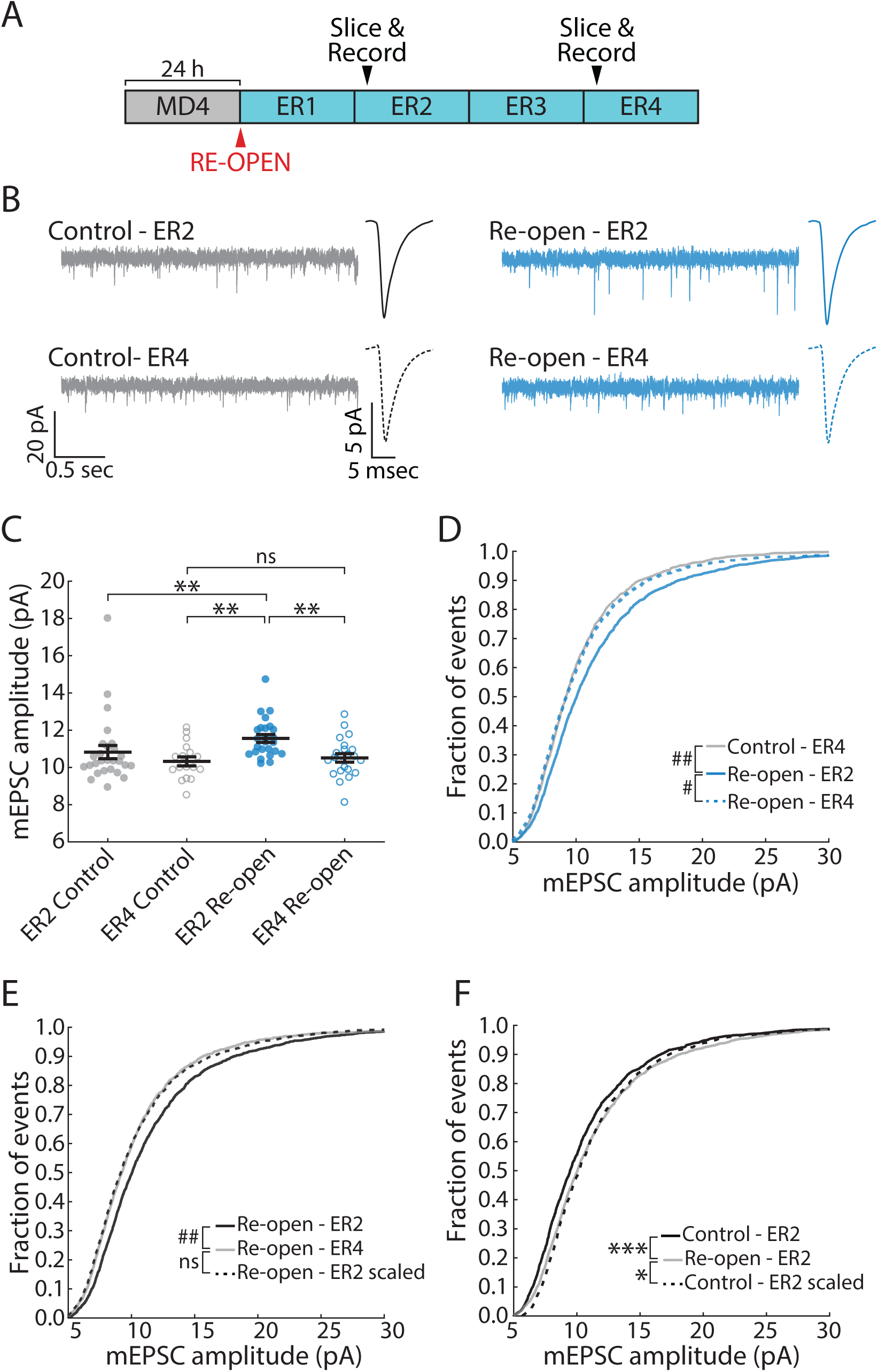
Downward FR recovery is associated with synaptic scaling down. **A**. Schematic of experimental timeline. Slices were taken 24 hr (ER2) or 72 hr (ER4) after ER. **B**. Example recordings of mEPSCs in L2/3 pyramidal neurons in V1. Average peak-aligned mEPSC waveforms for each condition are also shown. **C**. Average mEPSC amplitude in each condition. Each dot represents one cell. Black lines represent mean ± SEM. Kruskal-Wallis test (p < 0.001), Control ER2 n = 27, Control ER4 n = 17, Re-open ER2 n = 25, ER4 n = 22, ** p < 0.01. **D**. Cumulative distribution of amplitudes of mEPSC events. Kuiper test with Bonferroni correction, # p < 10^−5^, ## p < 10^−6^. **E**. Cumulative distribution of mEPSC amplitudes. Black dotted line represents Re-open ER2 distribution scaled according to the linear function *f*(*x*) = 0.877*x* + 0.318. Kuiper test, ## p < 10^−6^. **F**. Same as in E, but comparing the amplitude distribution at ER2 in Control vs Re-open conditions. The Control ER2 distribution has been scaled according to the linear function *f*(*x*) = 0.999*x* + 0.692. Kuiper test, * p =0.0377, *** p < 0.001. See Figure S4 for details of linear fit plots.

Homeostatic synaptic scaling affects mEPSC amplitudes in a uniform manner (Turrigiano et al., 1998). To test whether the change in mEPSC amplitude between ER2 and ER4 was consistent with synaptic scaling down, we plotted ranked re-opened ER2 amplitudes against ranked ER4 re-opened amplitudes and fit a linear function to the resulting plot (Figure S4). We then scaled the ER2 distribution using this function (Figure 3E). While the unscaled CDFs are significantly smaller on ER4 than ER2 (*p* < 10^−6^), the scaled-down ER2 CDF was almost identical to the ER4 CDF, and the two are statistically indistinguishable (*p* = 0.575), consistent with synaptic scaling. We next asked whether the potentiation of synaptic strength on ER2 was also consistent with synaptic scaling; in this case, we found that the scaled control CDF did not recapitulate the re-opened ER2 CDF and remained statistically distinct (*p* = 0.038, see also Figure S4), indicating a non-uniform change in strength across synapses, consistent with a non-homeostatic form of synaptic plasticity.

Taken together with the ability of NMDAR antagonists to block ER-induced potentiation of firing but not the homeostatic restoration of firing, these data support the conclusion that a Hebbian, LTP-like mechanism contributes to FR potentiation following ER, while synaptic scaling down contributes to downward firing rate homeostasis.

### Downward firing rate homeostasis is gated by sleep and inhibited by wake

We have previously shown that upward FRH is expressed only during periods of wake (Hengen et al., 2016), but it is unknown if homeostatic changes in the downward direction are regulated in this same way. We took advantage of our ability to record continuously from neurons during downward firing rate homeostasis while animals naturally cycle between periods of sleep and wake to investigate this. Animals’ behavioral state was classified using a supervised learning algorithm using a combination of local field potential (LFP), electromyogram (EMG) and video analysis (Figure S5). We first examined sleep- or wake-dense periods (2.5 hours with at least 70% sleep or wake; Figure 4A) over the 36-hour period when neuronal FRs on average are decreasing. Consistent with our previous results (Hengen et al., 2016), sleep and wake had no impact on FR in the control hemisphere. In striking contrast, in the ER hemisphere decreases in FR occurred exclusively during sleep-dense periods (Figure 4B). Results remained similar when key analysis parameters (such as window size and density percent threshold) were changed (Figure S6). The activity of most neurons in the re-opened hemisphere decreased across sleep-dense periods, but not during periods of wake (Figure 4C).

**Figure 4:**
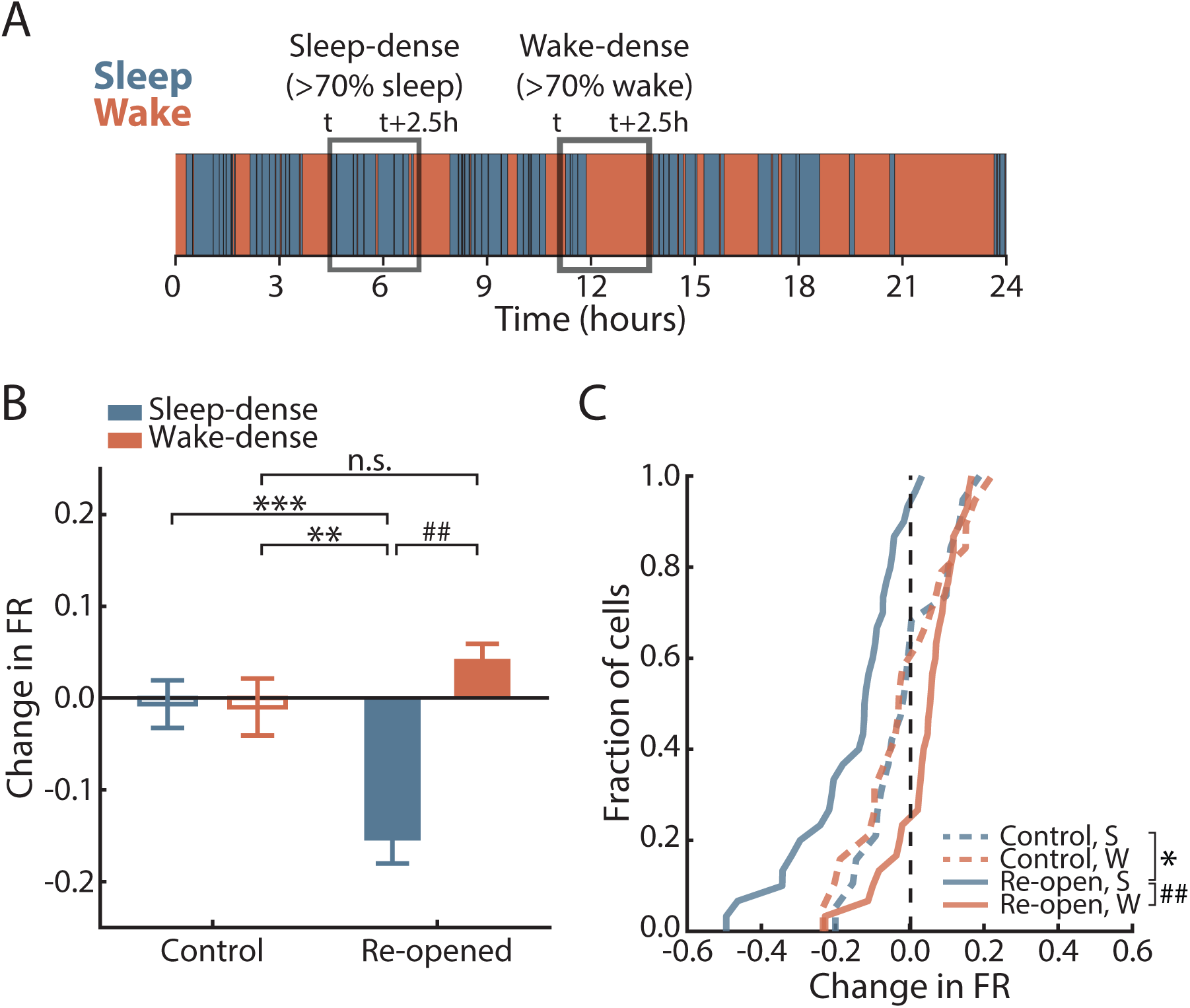
Downward FRH occurs during sleep-dense but not wake-dense periods. **A**. Schematic of state-dense analysis. **B**. Average change in FR over sleep- or wake-dense windows in Control and Re-opened hemispheres. Bars represent mean ± SEM. Kruskal-Wallis test (p < 10^−6^) with Tukey-Kramer post-hoc, Control Sleep-dense (S) n = 19 neurons, 13 windows; Control Wake-dense (W) n = 19 neurons, 10 windows; Re-open S n = 30 neurons, 12 windows; Re-open W n = 30 neurons, 11 windows; ** p = 0.0056, *** p = 0.0035, ## p < 10^−6^. This same result was obtained when using different analysis parameters (see Figure S6 for details). **C**. Cumulative distribution function of mean change in FR across S-dense or W-dense windows for all recorded cells. Black dashed line indicates no change. Two-sample Kolmogorov-Smirnov test with Bonferroni correction, Control neurons n = 19, Re-opened neurons n = 30, * p = 0.0258, ## p < 10^−6^.

We further classified behavior into four vigilance states: rapid eye movement sleep (REM), non-REM sleep (NREM), quiet wake (QW) and active wake (AW). Comparisons of mean FRs across these states are complicated by small but consistent differences in firing between them (Figure S5). We therefore used an approach based on an analysis of extended sleep or wake (defined as at least 30 min of sleep/wake without interruptions greater than 1 min; see also Miyawaki and Diba, 2016). In each of these periods, we measured the mean FR of neurons in a given state (e.g. NREM, for extended sleep). This allowed us to compare FRs in one state as a function of duration of time in that state (Figure 5A, B). We plotted the mean FR of each recorded neuron in the behavioral state of interest as a function of the time from the start of the extended sleep/wake episode, z-scored to the mean of the whole episode. In the re-opened hemispheres, there was a significant negative correlation between neuronal activity in NREM and time from the start of extended sleep (*r* = −0.193, *p* < 10^−30^), and a similar pattern in REM (*r* = −0.125, *p* < 10^−10^; Figure 5A). These correlations were absent in both NREM and REM in control neurons (Figure 5C), as well as in re-opened neurons during both wake states (Figure 5E), indicating that the decrease in firing was specific to extended sleep episodes in the re-opened hemisphere. Corroborating these results, we found a decrease in FR between the first and last NREM and REM episodes only in the re-opened hemisphere (Figure 5B). This decrease was absent in the other two conditions (Figure 5D, F).

**Figure 5:**
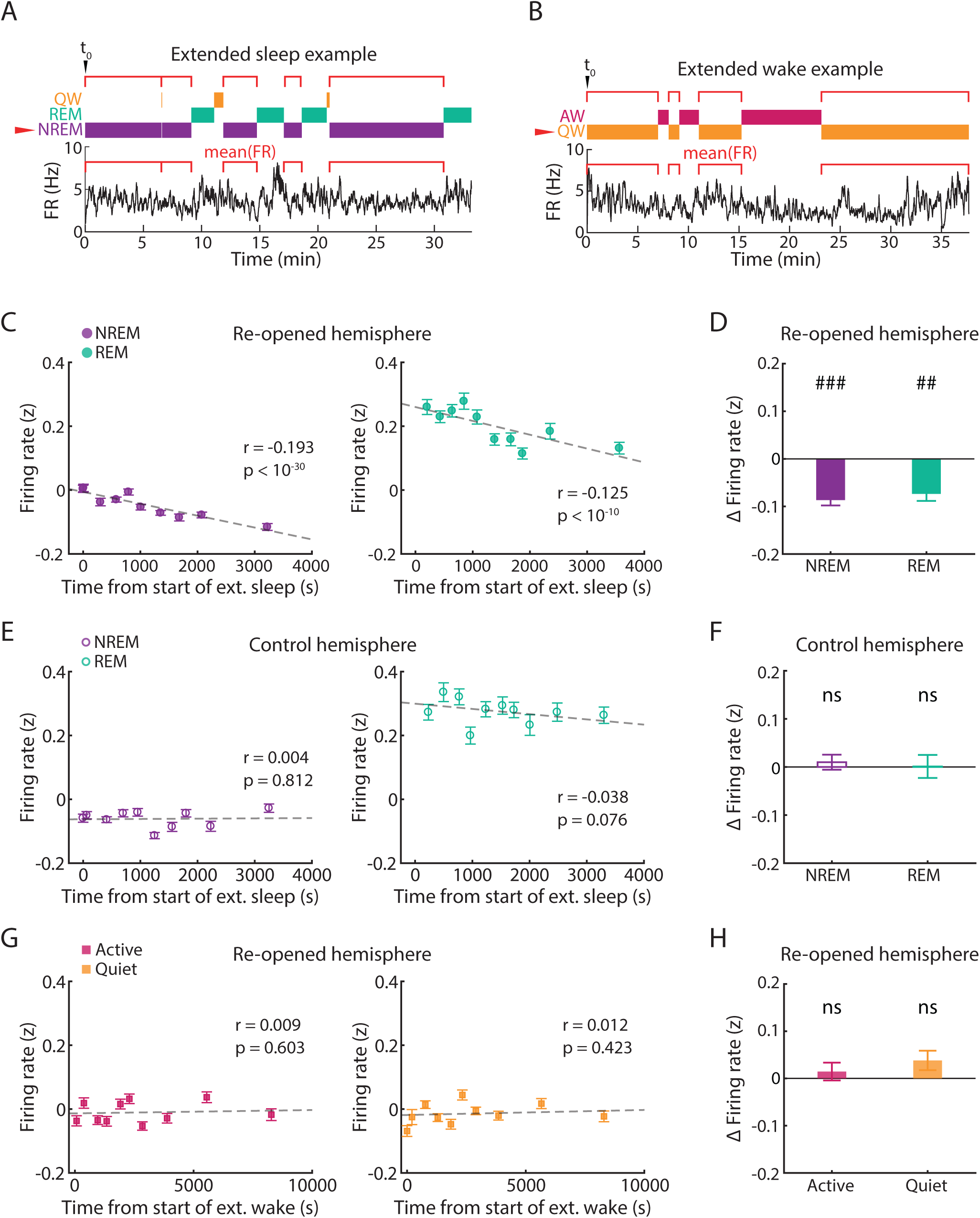
Downward FRH is gated by sleep and inhibited by wake. **A**. Schematic of extended sleep analysis for one neuron. Individual epochs of a given state (NREM in this example) were found within an extended sleep episode, and the mean FR of the neuron was calculated in each one. The values for each epoch were then plotted against the start time of that epoch, aligned to the start of the whole extended sleep episode (t0). **B**. Same as in A, but showing an example extended wake episode (same neuron as in A). **C**. Correlation between FR in NREM (left) and REM (right) and time from start of extended sleep, in the re-opened hemisphere. Data are grouped in 10 groups of equal size for visualization (dots show mean ± SEM for each group). Pearson *r* and associated p-values were computed on the ungrouped data (n = 3540 data points). FRs are z-scored to the whole extended sleep episode. **D**. Difference in FR between the last and first NREM (left) or REM (right) epoch within an extended sleep episode, averaged across all extended sleep episodes. Bars show mean ± SEM. One-sample t-test comparing to mean = 0, n = 62 episodes, ### p < 10^−13^, ## p < 10^−6^. **E, F**. As in C and D, but for active and quiet wake in the re-opened hemisphere (n = 3393 data points). In F, one-sample t-test comparing to mean = 0, n = 49 episodes. **G, H**. As in C and D, for NREM and REM in the control hemisphere (n = 2933 data points). In H, one-sample t-test comparing to mean = 0, n = 47 episodes.

Taken together these results show that downward firing rate homeostasis is happening exclusively during periods of sleep. Further, that control neurons show no decrease in firing during sleep demonstrates that sleep does not constitutively drive changes in firing rate, but rather enables the expression of downward firing rate homeostasis.

### Non-homeostatic FR changes happen independently of animals’ behavioral state

MD initially depresses firing rates over the first 2d of MD, via LTD at thalamocortical synapses and additional changes at intracortical synapses (Heynen et al., 2003; Maffei et al., 2006; Miska et al., 2018). This gives us the opportunity to test whether sleep and wake states also gate this non-homeostatic form of plasticity. We began by observing that during early MD, neuronal firing rates are stable for a variable period of time before starting to decline. Using an automated algorithm to detect the start of the drop for each neuron, we found that the drop happened quickly (over 6-12 hrs) in individual neurons (Figure 6A, B), but that the timing was variable; all our recorded regular spiking neurons could be classified as “early drop” neurons where most of the drop occurred during the first 12h light period after MD (starting 12h after MD; Figure 6A, C), or “late drop” neurons where most of the drop occurred during the second 12h light period after MD (starting 36h after the procedure; Figure 6B, D). When we analyzed the relationship between sleep/wake behavior and the drop in neuronal FRs during the first or second 12h light periods (for early and late drop neurons, respectively) we found that in neither group was there a bias towards wake or sleep. The magnitude of the decrease in FR was similar across both behavioral states (Figure 6E), and it was not correlated with the amount of time spent asleep (Figure 6F). Thus, we conclude that sleep specifically enables a reduction in firing rate driven by homeostatic plasticity.

**Figure 6:**
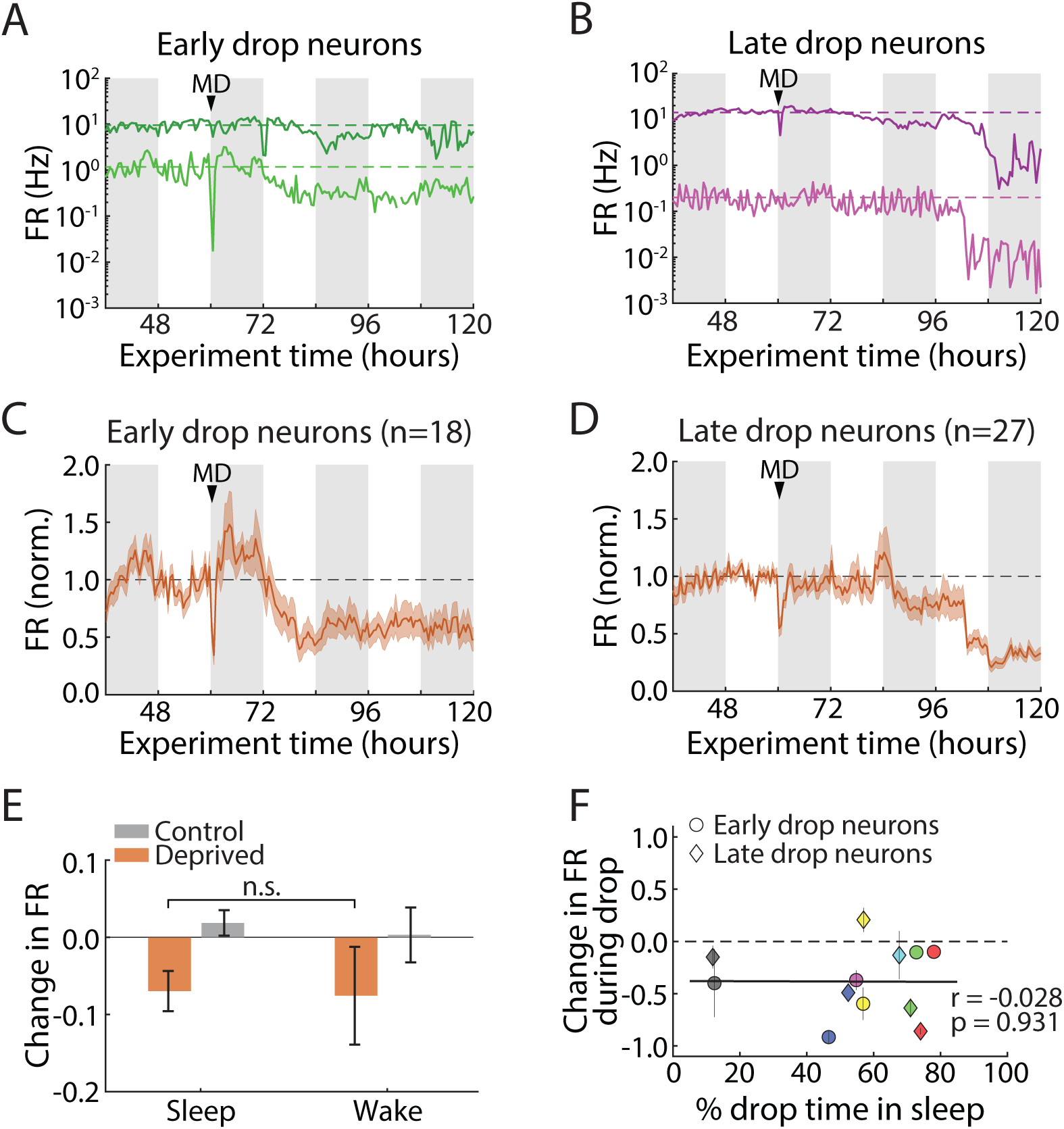
Non-homeostatic FR changes happen independently of animals’ behavioral state. **A**. Two examples of neurons whose activity begins decreasing in the first light period after MD. Dashed lines indicate baseline FR for each neuron. White/gray bars in background indicate 12 hours of light/dark. **B**. As above, but for two example neurons for which the drop in FR occurs in the second light period after MD. **C**. Average baseline-normalized FR of all “early drop” neurons, i.e. for which drop began in the first light period after MD. Dashed line indicates baseline FR. **D**. As in C, but for “late drop” neurons. **E**. Change in FR across sleep- or wake-dense epochs for all neurons in their respective 12-hour drop period (1^st^ light period after MD for “early drop”, 2^nd^ light period after MD for “late drop”). Control S-dense n = 38 epochs, W-dense n = 12 epochs; Deprived S-dense = 38 epochs, W-dense n = 26 epochs. **F**. Correlation between change in FR during drop and percent time spent asleep in the same period. Each data point represents the average change in FR across early drop (circles) or late drop (diamonds) neurons, and each color represents a different animal (n = 6 animals). Percent time asleep is calculated per animal in the early or late drop 12-hour period. Black dashed line represents no change. Black solid line shows linear fit to the data.

## DISCUSSION

It has been theorized that bidirectional stabilization of firing rates around an individual set point is a critical feature of many neuronal networks, that allows developmental or experience-dependent synaptic changes to refine network architecture without fatally destabilizing activity (Abbott and Nelson, 2000; Marder and Prinz, 2002; Tetzlaff et al., 2011; Turrigiano, 2012; Litwin-Kumar and Doiron, 2014). While upward firing rate homeostasis is known to occur *in vivo*, whether this process is indeed bidirectional, and whether upward and downward homeostasis share mechanistic features, have been open questions. Here we used an MD followed by ER paradigm to potentiate firing in V1, then analyzed the behavior of individual neurons over time in freely behaving animals. MD-ER produced a ∼2-fold potentiation of firing that was NMDAR-dependent and accompanied by non-uniform changes in synaptic strengths. This was followed by downward FRH, which slowly returned individual firing rates close to their initial values. This homeostatic recovery of firing was independent of NMDAR signaling, and was accompanied by a scaling down of synaptic strengths, suggesting it is driven at least in part by homeostatic synaptic scaling. Strikingly, we found that downward FRH is promoted by sleep and inhibited by wake, exactly the opposite of upward firing rate homeostasis (Hengen et al., 2016). This does not reflect a general role for sleep in all forms of synaptic depression, as the early phase of MD (driven by LTD-like mechanisms) unfolded independently of sleep and wake states. Our data show that the role of sleep/wake states in promoting circuit plasticity is quite nuanced, and that the induction of upward and downward FRH are segregated by behavioral state.

While sensory deprivation is a venerable paradigm for inducing neocortical plasticity (Espinosa and Stryker, 2012; Gainey and Feldman, 2017), there has been a dearth of approaches for increasing, rather than decreasing, neocortical firing to study downward homeostatic plasticity. Prolonged dark exposure is one paradigm that has been proposed to induce homeostatic plasticity within V1 (Goel and Lee, 2007), but more recent work has shown that changes in mEPSC amplitude during dark exposure unfold via metaplastic rules that are distinct from those that induce synaptic scaling (Bridi et al., 2018). Dark exposure does not impact firing rates in V1 over the first day or so (Torrado Pacheco et al., 2019) and so would not be expected to induce synaptic scaling, and after longer dark exposure there is disagreement as to whether firing rates increase (Bridi et al., 2018) or decrease (Torrado Pacheco et al., 2019); in both studies these effects were subtle. Even light re-exposure after 60 hr in the dark only elevates firing for ∼20 minutes, not long enough to trigger slow forms of homeostatic plasticity (Torrado Pacheco et al., 2019). In contrast, MD followed by ER induces a robust and long-lasting increase in firing in V1. This potentiation of firing is not instantaneous but develops over several hours, and is blocked by NMDAR antagonists, suggesting (consistent with previous work, Toyoizumi et al., 2014) that it is a consequence of synaptic plasticity induced by the restoration of correlated visual drive through the previously closed eye.

Consistent with our findings, CPP has been shown to block electrically induced LTP and LTD *in vivo* in neocortex and hippocampus (Trepel and Racine, 1998; Villarreal et al., 2002; Medvedev et al., 2010) and to block the early, Hebbian phase of MD-induced ocular dominance (OD) plasticity in V1, but not the late phase which depends on upward homeostatic plasticity (Sato and Stryker, 2008; Toyoizumi et al., 2014). There has been some disagreement over whether blocking NMDAR impacts visual function, but several recent studies have shown that CPP injections in critical period rodents at the dose used here do not impact visual acuity or other aspects of visual function (Sato and Stryker, 2008; Toyoizumi et al., 2014). We see only a small and short-lived effect of CPP injections on FRs under control conditions (Figure S3), much shorter than ER-induced potentiation (Figure 1D). Taking these considerations together, the most likely explanation for the ability of CPP to block potentiation of firing rates is that it prevents Hebbian potentiation caused by the reinstatement of correlated visual drive to a cortical circuit that is in a high-gain state following upward firing rate homeostasis (Lambo and Turrigiano, 2013 Hengen et al., 2013), an interpretation that is consistent with simulations of Hebbian and homeostatic interactions within V1m (Toyoizumi et al., 2014).

Similar to the homeostatic phase of OD plasticity (Toyoizumi et al., 2014), downward FRH is independent of NMDAR signaling. This is most consistent with the induction of homeostatic forms of plasticity that do not rely on calcium influx through NMDAR, such as synaptic scaling. *In vitro* studies have revealed a dependence of scaling down on NMDAR in younger neurons (Hou et al., 2011), but not in cultured slices or older neurons (> 3 weeks *in vitro*), where scaling down is instead dependent on activation of L-type calcium channels (Goold and Nicoll, 2010; Siddoway et al., 2013). Two other *in vitro* studies reported that bicuculline-induced scaling could be attenuated by blocking NMDARs (Lissin et al., 1998; Qiu et al., 2012), but neither of these controlled for changes in firing rate due to NMDAR-blockade in disinhibited cultures, which can explain this effect (Leslie et al., 2001). Further, downward FRH after ER was associated with a global scaling down of synaptic strengths in V1m. Taken together with the lack of effect of NMDAR blockade, this supports a model in which downward FRH is driven (at least in part) by homeostatic downscaling of excitatory synaptic strengths.

The process of downward FRH revealed here bears a number of similarities to upward FRH (Hengen et al., 2016). They both unfold slowly over ∼2 days, are accompanied by synaptic scaling of excitatory synapses in the correct direction to contribute to restoration of firing, and bring individual neuronal firing rates back to an individual firing rate set point. It was thus surprising to find that they have a diametrically opposite dependence upon behavioral state: upward FRH occurs during active waking, while downward FRH occurs during sleep. These findings show that upward and downward homeostatic compensation do not operate simultaneously within neuronal circuits, and add complexity to the influential idea that the function of sleep is to provide a window of opportunity for homeostatic plasticity (Wang et al., 2011).

The synaptic homeostasis hypothesis (SHY) proposes that wakefulness drives learning-related increases in synaptic strengths and FRs, while sleep renormalizes activity by down-regulating synaptic strengths, a process dependent on activity patterns during NREM sleep (Tononi and Cirelli, 2014). A combination of structural, electrophysiological and molecular evidence has been put forward in support of SHY (de Vivo et al., 2017; Vyazovskiy et al., 2008; Vyazovskiy et al., 2009; Liu et al., 2010; Diering et al., 2017), but contradictory data have also been reported (Yang et al., 2014; Chauvette et al., 2012; Aton et al., 2014; Durkin and Aton, 2016), and in some cases the interpretation of these findings has been questioned (Frank, 2012; Frank and Cantera, 2014; Timofeev and Chauvette, 2017; Puentes-Mestril and Aton, 2017). In particular, it has been unclear whether sleep and wake cause global oscillations in firing rates under baseline conditions, when animals have not experienced a dramatic plasticity or learning event. Such oscillations have been observed in the hippocampus and frontal cortex, but were not observed uniformly across the neuronal population (Miyawaki and Diba, 2016; Watson et al., 2016). In the absence of plasticity induction, we observe stable firing during even relatively long, consolidated sleep/wake states in V1, both here and in previous datasets (Hengen et al., 2016), indicating that in V1 sleep and wake to not drive global, brain-wide changes in excitability under basal conditions. Further, we find that downward firing rate changes driven by Hebbian plasticity during early MD unfold independently of sleep and wake states. Thus it is not the case that all decreases in synaptic strength and firing preferentially occur during sleep.

Our results paint a nuanced and complex picture of the impact of sleep and wake on neuronal plasticity. The impact of sleep on FR changes is not uniform, but depends on the particular forms of synaptic plasticity activated, an idea which may help explain the diversity of outcomes in similar experiments designed to test the role of sleep in the induction of plasticity (Raven et al., 2018; Frank and Seibt, 2018). While our data argue against a constitutive role of sleep in reducing circuit excitability (Figure 5 and see Hengen et al., 2016), we do find that when it is induced through manipulations of experience, downward FRH (and presumably the underlying downscaling of synapses) occurs preferentially during sleep. This finding is consistent with a recent study showing that sleep deprivation interferes with molecular signaling cascades that are important for scaling down (Diering et al., 2017). The most parsimonious explanation for our data is that – rather than constitutively inducing downscaling – sleep is permissive for the expression of downward homeostatic plasticity when it is induced by perturbations to the circuit.

A surprising finding of this study is that upward and downward FRH are gated by distinct sleep/wake states. What could be the mechanism by which this is achieved, and what is the purpose of this behavioral state segregation? One promising lead may be the starkly contrasting neuromodulatory environments neocortical neurons are exposed to during sleep and wake (Lee and Dan, 2012). Neuromodulators may regulate plasticity mechanisms by modulating neuronal activity (Goard and Dan, 2009; Herrero et al., 2008), or by directly acting on signaling pathways involved in plasticity induction (Diering et al., 2017). The molecular pathways that regulate synaptic scaling up and down are mostly divergent, but do show some overlap (Turrigiano, 2017; Styr et al., 2019). One possible reason for the temporal segregation of upward and downward homeostasis could be to reduce interference or saturation of the molecular pathways that mediate homeostatic plasticity. A complementary possibility is that behavioral state gating ensures strong unidirectional homeostatic compensation for defined periods of time; this may have benefits in terms of computation and learning, by providing strong compensation when it is needed most, and perhaps by allowing unopposed Hebbian changes during limited windows of time.

## ACKNOWLEDGEMENTS

Funding sources: R01-EY025623 (G.G.T.). Computational resources were provided by NSF XSEDE computing resources (MCB090163) and the Brandeis HPCC which is partially supported by DMR-1420382.

## AUTHOR CONTRIBUTIONS

Conceptualization, A.T.P., G.G.T.; Methodology, A.T.P., J.B., G.G.T.; Software, A.T.P., J.B.; Formal Analysis, A.T.P., J.B.; Investigation, A.T.P.; Writing, A.T.P., G.G.T.; Visualization, A.T.P., J.B.; Data Curation, A.T.P.; Supervision, G.G.T.; Funding Acquisition, G.G.T.

## STAR METHODS

### Contact for Reagent and Resource Sharing

Further information and requests for resources and reagents should be directed to the lead contact, Gina G. Turrigiano.

### Experimental Model and Subject Details

All animal care as well as surgical and experimental procedures were approved by the Animal Care and Use Committee (IACUC) of Brandeis University and complied with the guidelines of the National Institutes of Health. All experiments were performed on Long-Evans rats of both sexes (Charles River Laboratories, Wilmington, MA, USA; strain code: 006) aged P21-P35. Timed pregnant female Long-Evans rats were obtained and housed on a 12h/12h light/dark cycle with free access to food and water. For *in vivo* electrophysiology experiments, 2 subjects from the same litter were weaned at post-natal day 21 (P21) for electrode implant surgeries and then housed together in a satellite facility. For slice physiology, rats were weaned at P21, housed in the main animal facility along with littermates, and returned there after every surgical procedure.

### Method Details

#### Surgical procedures

##### Electrode implants

Rats were implanted with electrode arrays as described previously (Hengen et al., 2013). Briefly, custom 16-channel tungsten wire (33 μm tip diameter, Tucker-Davis Technologies, TDT, Alachua, FL, USA) arrays were implanted bilaterally in V1m. Anesthesia was induced with an intraperitoneal injection of ketamine/xylazine/acepromazine (KXA) cocktail (70 mg/kg ketamine; 3.5 mg/kg xylazine; 0.7 mg/kg acepromazine) and maintained with isoflurane (1.0% – 2.0% concentration in air) delivered via an anesthesia system with integrated digital vaporizer (Somnosuite, Kent Scientific, Torrington, CT, USA) through a stereotaxic head holder (Model 923-B with Model 1924-C-11.5 mask, Kopf Instruments, Tujunga, CA, USA). The skull was exposed, cleaned with hydrogen peroxide, and any bleeding spots were lightly cauterized. Three small holes were drilled in the bone, one above the cerebellum and two above motor/somatosensory cortex, and miniature machine screws (Antrin Inc., Fallbrook, CA, USA) were inserted in each. A craniotomy was drilled above V1m on the left hemisphere and the *dura mater* was pulled back using a 25G needle. The electrode array was then slowly lowered into the brain and the exposed craniotomy was covered with a silicone elastomer (Kwik-Cast, World Precision Instruments, Sarasota, FL, USA). The array was secured using dental cement, then grounded to both front screws using steel wire and soldering paste. The same procedure was repeated for the right hemisphere, and both arrays were secured to the screws and bone surface using dental cement. Total headcap weight was approximately 2g. Finally, two braided steel wires were implanted deep in the nuchal muscle for EMG recordings.

##### Monocular deprivation

For MD, rats were briefly (∼20 sec) administered 4% isoflurane, then transferred to a heating pad and placed in a nose cone that delivered 1.0 – 3.0% isoflurane in air. Ophthalmic ointment was applied to the eye not being sutured to prevent desiccation. The other eyelid was cleaned 3 times with betadine followed by flushing with sterile saline. Lidocaine cream was applied to the eyelid. The bottom and top part of the eyelid were then sutured together using 6-0 nylon or polyester sutures (4 mattress sutures). The sutured eye was covered in antibiotic ointment and lidocaine, and analgesic (meloxicam, 1 mg/kg) was administered subcutaneously. The lidocaine, antibiotic and analgesic were given again 24 hours after surgery. Sutures were checked daily and animals were excluded if sutures were not intact at the time of ER.

##### Eye re-opening

Rats were anesthetized as for MD. Ophthalmic ointment was applied to the non-sutured eye. The sutured eye was cleaned with betadine and saline thrice. Sutures were then carefully cut with fine-tipped surgical scissors, and removed using small forceps. The re-opened eye was flushed with saline until it was free of any extraneous tissue and looked clean. Ophthalmic antibiotic ointment was applied, and the animal returned to the cage. Occasionally (3 animals), the eye was found to have developed infection or a cataract during the lid suture period. These animals were excluded from the study.

#### Continuous single-unit recordings in freely behaving animals

Following electrode implant surgery rats were allowed to recover for 2 days in separate cages with *ad lib* access to food and water. During this time animals were handled twice daily by experimenters or assistants, placed in the same cage for 10 minutes twice daily to allow for social interaction, and given treats (Froot Loops). The evening before the recording started animals were transferred to a clear plexiglass cage of dimensions 18”x12”x18” (length, depth, height) and separated into two arenas by a clear plastic divider with 1” holes to allow for tactile and olfactory interaction between siblings while preventing aggressive play and jostling of headcaps. Animals were kept on a 12h/12h light/dark cycle in a temperature- and humidity-controlled room (lights on 7:30am, 21°C, 25-55% humidity). The arrays were connected to TDT commutators via ZIF-clip headstages to allow animals to behave freely. Data were recorded continuously for 9-12 days (up to ∼250 hours). Animals were only disconnected for MD and ER procedures (∼20 min each animal). Data were acquired at 25 kHz, digitized and streamed to disk online using a TDT Neurophysiology Workstation and Data Streamer. Spike extraction, clustering and sorting were done using custom Matlab and Python code (see below).

#### Automated spike extraction, *clustering and sorting*

Spike extraction, clustering and sorting were done as previously described (Hengen et al., 2016; Torrado Pacheco et al., 2019). Spikes were detected in the raw signal as threshold crossings (−4 standard deviations from mean signal) and re-sampled at 3x the original rate. Principal component analysis (PCA) was done on all spikes from each channel, and the first four principal components were used for clustering (Harris et al., 2000). A random forest classifier implemented in python was used to classify spikes according to a model built on a dataset of 1200 clusters manually scored by expert observers. A set of 19 features, including ISI contamination (% of ISIs < 3 msec), similarity to regular spiking unit (RSU) and fast-spiking unit (FS) waveform templates, amount of 60 Hz noise contamination and kinetics of the mean waveform. Cluster quality was also ensure by thresholding of L-ratio and isolation distance (Schmitzer-Torbert et al., 2005). Clusters were classified as noise, multi-unit or single-unit. Only single-unit clusters with a clear refractory period were used for FR analysis. We classified units as RSU or FS based on established criteria (mean waveform trough-to-peak and tail slope, Hengen et al., 2016). Only RSUs (putative excitatory neurons) were used for analysis.

Some neurons were lost during the recording, presumably due to electrode drift or gliosis. To establish “on” and “off” times for neurons, we used ISI contamination: when hourly % of ISIs < 3 msec was above 4%, unit was considered to be offline. Based on these “on” and “off” times, only units that were online for 80% of the 5-day period were used for analysis. Additionally, we used a stringent post-hoc bootstrap analysis (Figure S1) of daily average waveforms to discriminate between units that were followed continuously for 5 days versus multi-unit signal. Daily mean waveforms (WFs) for 5 days (MD4 to ER4) were computed for all putative single-units; then for each unit we randomly picked 3 of its WFs and mixed them with two other WFs from a randomly selected cell in the dataset. Based on these 5 WFs, calculated the daily mean-squared error (MSE) between them, and found the maximum MSE for shuffled unit. This process was repeated 1000 times per cell, to obtain a distribution of random unit maximum MSEs. We chose maximum as opposed to summed or average MSE to increase stringency: this method results in a high MSE for a WF that is stable for 4 days but changes in the last day, for example. To obtain an MSE threshold, we chose the lower bound of the 95% confidence interval for the mean of the distribution of random unit maximum MSEs. Putative units with maximum MSE across days greater than this value were excluded from analysis. The resulting real units had very similar WFs across days and low maximum MSE (Figure S1).

#### Transcardial perfusions

Animals were deeply anesthetized with a double dose of KXA (see above for full dose). The heart was exposed and a 21G needle inserted in the left ventricle. After cutting a small hole in the right atrium, 0.9% saline was perfused through the circulatory system using a peristaltic pump for 5-7 minutes. The perfusion was switched to 3.7% paraformaldehyde (PFA) for another 5-7 minutes. The brain was extracted taking care not to damage the electrode insertion site, and preserved in a solution of 3.7% PFA and 30% sucrose for at least 7 days for cryo-protection.

#### Histology

Fixed brains were blocked using a razor blade and mounted on a freezing microtome platform kept cold by dry ice. Brains were embedded in O.C.T. compound (Tissue-Tek, Sakura, Japan) and 60 µm thick sections were taken and placed in phosphate-buffered saline (PBS) overnight. Slices were then stained with cresyl violet to dye the Nissl substance in neurons. Stained sections were mounted and coverslipped, then imaged at 4x or 10x on a digital microscope (Keyence, Belgium) to confirm the location of each electrode wire.

#### CPP injections

All CPP injections were done on animals that were undergoing chronic electrophysiological recordings. Animals were not unplugged for this procedure, unless eye re-opening surgery was also performed. After weighing, animals were administered a 15 mg/kg dose of (RS)-CPP (Tocris, Bio-Techne corp., Minneapolis, MN, USA) dissolved in bacteriostatic 0.9% saline subcutaneously.

The CPP solutions were prepared on the day of injection, and re-used the next day when applicable (storing overnight at 4°C).

#### Slice electrophysiology

#### Standard ACSF (in mM)

126 NaCl, 25 NaHCO3, 3 KCl, 2 CaCl2, 2 MgSO4, 1 NaH2PO4, 0.5 Na-Ascorbate, with dextrose added to bring osmolarity to 310-315 mOsm, and titrated with HCl to bring pH to 7.35.

#### TTX-ACSF

standard ACSF with added tetrodotoxin (TTX), 0.2 µM.

#### Choline solution (in mM)

110 Choline-Cl, 25 NaHCO3, 11.6 Na-Ascorbate, 7 MgCl2, 3.1 Na-Pyruvate, 2.5 KCl, 1.25 NaH2PO4, and 0.5 CaCl2, with dextrose added to bring osmolarity to 315 mOsm, and titrated with HCl to bring pH to 7.35.

#### K-gluconate internal solution (in mM)

100 K-gluconate, 10 KCl, 10 HEPES, 5.37 Biocytin, 10 Na-Phosphocreatine, 4 Mg-ATP, and 0.3 Na-GTP, with sucrose added to bring osmolarity to 295 mOsm and KOH added to bring pH to 7.35.

##### Acute brain slice preparation

Coronal brain slices (300 µm) containing V1 from both hemispheres were prepared using a procedure similar to one used in previous studies (Miska et al., 2018). Animals were placed in a sealed container with 4% isoflurane in air and deeply anesthetized. They were then quickly decapitated and the front part of the brain (excluding the cerebellum and part of the brainstem) was extracted within 60 sec and placed in cold (∼1°C) carbogenated (95% O2, 5% CO2) TTX-ACSF for 4 min. Once the brain was cold and firm, it was cut coronally through frontal cortex to obtain a flat mounting surface, and mounted to a slicing chamber using cyanoacrylate adhesive. Slices were immediately cut on a vibratome (Leica VT1000S, Diegem, Belgium) in cold carbogenated TTX-ACSF. Immediately after cutting, each slice was transferred to an incubation chamber placed in warm (34°C) continuously carbogenated choline solution for protective recovery. After 10 min, slices were transferred to warm (34°C) continuously carbogenated TTX-ACSF for 40 min. They were then removed from the incubator, placed in room temperature TTX-ACSF and allowed to return to room temperature before recording. TTX-ACSF was used throughout to prevent additional plasticity due to activity in the slice after cutting. Slices were used for recordings for up to 6 post-slicing.

##### mEPSC recordings

Borosilicate glass pipettes were pulled on a Sutter P-97 Micropipette puller. Pipettes were used if they had tip resistances ranging from 4-6 MΩ, and filled with K-gluconate internal solution. V1m was identified in acute slices using the rat brain atlas (Paxinos and Watson, 1998) based on morphology of the hippocampus and white matter, and a high-contrast band corresponding to layer 4 (L4). Pyramidal L2/3 neurons were identified by their position (dorsal to L4), teardrop-shaped soma and presence of an apical dendrite. This was confirmed by post-hoc reconstruction of biocytin fills. On any given recording day cells were patched from both re-opened and control hemispheres. All recordings were performed in submerged slices continuously perfused with carbogenated ACSF at 32°C. Cells were visualized on an upright microscope (Olympus BX51WI) using a 10x air (NA 1.13) or 40x water-immersion (NA 0.8) objective and an infrared CCD camera. Cells were patched using pipettes filled as above and with a chlorided silver electrode. Data were low-pass filtered at 5 kHz acquired at 10 kHz using a National Instruments Data Acquisition Board (DAQ, National Instruments, Woburn, MA, USA) and custom Matlab software. All post-hoc analyses were done using in-house software written in Matlab. For mEPSC recordings, TTX-ACSF with added AP-5 (50 µM) and picrotoxin (25 µM) was used to block action potentials and NMDA and GABA currents, and isolate AMPA currents. Neurons were held in voltage clamp at −70 mV while at least 20 traces (10 sec duration) were recorded at 10x gain. Neurons were excluded from analysis if series resistance was above 25 MΩ, if resting membrane potential (V_m_) was above −60 mV, or if V_m_ changed by more than 10% during trace acquisition. For event detection, 3 traces with stable baseline were selected. In order to both reliably detect mEPSC events above noise and to limit bias in selection, we used an in-house program written in MATLAB that employs a semi-automated template-based detection method contained in a GUI. In brief, the program first filters the raw current traces and then applies a canonical mEPSC event shaped template to detect regions of best fit. Multiple tunable parameters for template threshold and event kinetics that aid in detection were optimized and then chosen to stay constant for all analyses. Putative events are then analyzed for hallmark features of mEPSC (risetime kinetics, decay time, minimum amplitude cutoffs, etc.). Finally, the results of the automated detection are reviewed with minor manual revisions made (<5%) for the inclusion/exclusion of events. We performed all of this analysis (including the manual revisions) blind to experimental conditions. Neurons were excluded from analysis post-hoc if they were determined to be non-pyramidal based upon biocytin reconstruction.

#### Semi-automated behavioral state scoring

Local field potentials (LFPs) were extracted from the raw *in vivo* electrophysiology traces by low-pass filtering data from 3 channels at 300 Hz, resampling at 200 Hz, and averaging. We computed power spectral densities using Matlab’s fast-fourier transform (FFT) implementation in 10 sec bins and frequency bins from 0.3 to 15 Hz (0.1 Hz steps). Power in the delta (0.3 – 4 Hz) and theta (5 – 8 Hz) bands was calculated in each 10-sec bin as a fraction of total power. EMG signals from the two wires were averaged and resampled at 200 Hz. Animal movement was tracked in recorded videos (recoded using an infrared camera) using open-source software written in C++ (Video Blob Tracking, Open Source Instruments, Watertown, MA, USA; Github: https://github.com/OSI-INC/VBT) and modified in-house to suit our needs (adding video cropping, tracking in both light and dark conditions). For sleep classification, we built a custom graphical user interface (GUI) in Matlab. Data was presented to an observer in 1-hour blocks and scored in 10-sec bins. Scoring of the first 10 blocks was manual, based on previously published criteria (Hengen et al., 2016; Torrado Pacheco et al., 2019), and included 4 states: NREM sleep (high delta power, low theta power; low EMG and movement); REM sleep (low delta power; high theta power; lowest EMG and movement); quiet wake (low delta and theta; low EMG and movement); active wake (low delta and theta; higher EMG and movement). After block 10, a random forest classifier model (Python Scikit-Learn implementation) was built based on scoring of blocks 1-10. The features used in the model were power in the delta, theta and gamma (40 – 100 Hz) bands, theta-delta power ratio, z-scored EMG and movement signal. The model was used to classify new data (blocks 11 and above), and the result was displayed in the GUI. Trained human scorers were then able to check LFP power, EMG and movement traces, as well as view video recordings, to correct the classification. The model was updated with each new scored block until the training set reached 10,000 bins (limit set for efficiency and speed). Thereafter, the training set was continuously updated to contain the most recently coded 10,000 bins. The GUI also allowed experimenters to exclude a block for training (in case of corrupted data, or animal unplugging for surgery). This algorithm reached an OOB error of <10% and real error (i.e. % of bins that had to be corrected) of 1-4% in sleep-heavy blocks, and 5-15% in wake-heavy blocks, the main difficulty being distinguishing quiet vs active wake.

#### FR analyses

##### Estimation of neuronal FRs and change in FR

To obtain FRs estimates for individual RSUs from spike timestamps we computed spike counts in 60-second bins and applied a Gaussian kernel with σ = 300 seconds (Figures 1, 2, 4, 6), or we computed the spike rate in non-overlapping 1-second bins (Figure 5). To calculate mean FRs in 12-hour periods we took the average FR across all bins in that period (Figures 1, 2). FRs were normalized to baseline by dividing all FR bins by the mean FR in the chosen 12-hour baseline period (Figures 1, 2, 6). To calculate FR z-scores over an episode of extended sleep or wake (Figure 5) we computed the mean and SD of the non-normalized FR over that episode and applied the formula *Z*_*FR*_ = (*FR* − *μ*)/*σ*, where *μ* and *σ* are the mean and SD and FR is the non-normalized FR. To estimate the change between first and last epochs of a state in extended sleep or wake (Figure 5) we subtracted the mean z-scored FR of the first epoch from the last one. To estimate the changes in FR in sleep- and wake-dense windows we used the formula (*B* − *A*)/(*A* + *B*). For the result in Figure 4, we defined *A* and *B* as the mean FR in the first and last 15 minutes of the S- or W-dense window. For the MD-induced drop (Figure 6E) we defined *A* and *B* as the mean FR in the first and last 40% of the window. To estimate the total change in FR across the drop we used the same approach, but with *A* being the mean baseline FR and *B* being the mean FR in last 20% of the drop period (Figure 6F).

##### Estimating start of MD-induced drop

To estimate the start of the drop for each neuron in an unbiased way, we designed the following algorithm. A 2-hour window was stepped through the data in 15-minute increments and the neuron’s FR in that window was fit with a linear function to calculate the slope. Negative and positive slopes at least one s.d. from the mean slope were selected, and a kernel density estimate (KDE) was calculated separately for the positive and negative ones. The drop start time was identified as the first time that the difference between the negative and positive KDE went above 70%. This start time was then used to classify neurons as “early dorp” or “late drop” depending on whether it fell within the first or second light period after MD.

#### Bootstrap analysis of FR recovery

For the result in Figure S2, we used two separate bootstrap methods to analyze the FR recovery. Our starting distributions were the data in Figure 1G (right). We aimed to compare the baseline (MD4) distribution to the late ER (ER4) distribution to ask if the return to baseline activity was happening cell-autonomously. The % change from baseline for the real late ER distribution was (2% ± 13%) in Figure 1F. Could this result be obtained if instead of each cell returning to the observed value, we either shuffled the end points (i.e. keeping the resulting mean population FR constant, but shuffling neurons’ place in the distribution), or returned each neuron to a different value sampled at random from the late ER distribution? Both the shuffling and sampling analyses were done 10,000 times to obtain 10,000 bootstrap means and corresponding confidence intervals. The results in Figure S2C show that the value obtained in our experiments is outside the 99% confidence interval for both analyses. We also aimed to determine the range of variation of FR in the real data. In other words, how precisely do neurons return to their baseline FR? To do this we calculated the fraction of neurons returning to within X% of their baseline FR, where X was varied from 10% to 100%, in both our real data and for both bootstrap methods. We used the bootstrap distributions to obtain 99% CIs for the fraction of neurons at each % threshold value. We find that more than 50% of real neurons return to within 50% of their baseline FR, and the real value diverges from the bootstrap results at the 40% threshold (Figure S2D).

#### Sleep analyses

##### Sleep- and wake-dense windows

For the results in Figure 4, we scanned hours 192-240 of the experiment (ER2-ER4) in 15 min steps for 2.5 h periods of time where animals had been awake or asleep for at least 70% of the time. When a dense window was found the algorithm restarted scanning at the end of that window (i.e. windows were not double-counted). The FR change was calculated for each unit as *(B – A)/(B + A)*, where *A* and *B* represent the mean FR in the first and last 15 min of the window. We repeated this analysis with different parameters to confirm that our results were not spurious: similar results were obtained when changing the size of the window and the density % threshold (Figure S6).

##### Extended sleep and wake analyses

For the results in Figure 5, we scanned hours 192-240 of the experiment for periods of extended wake or sleep, defined as at least 30 min of wake or sleep without interruptions greater than 1 min. Only states that were 30 sec or longer were considered for this analysis. The FR for each cell was z-scored to the mean and s.d. of the FR for the whole extended sleep or wake episode.

### Quantification and Statistical Analysis

All data analysis was done using custom code written in Matlab. Values are reported in the text body as mean ± SEM. For statistical analyses n’s, p-values and the kind of test used are provided in the figure caption. Normality was assessed using an Anderson-Darling test (Matlab implementation), with α = 0.05. To compare means across groups for normally distributed data we used one-way ANOVA followed by Tukey-Kramer post-hoc for pairwise comparisons, or 2-sample t-tests to compare two groups only. For non-normally distributed data we used a Kruskal-Wallis test followed by Tukey-Kramer post-hoc for pairwise comparisons. For non-normal paired data we used Wilcoxon sign-rank tests followed by Bonferroni corrections for multiple comparisons. To compare cumulative distributions we used a two-sample Kolmogorov-Smirnov (K-S) test or a two-sample Kuiper test with Bonferroni corrections for multiple comparisons. Correlations strength and significance were estimated using Pearson’s *r*.

### Data and Code Availability

Data and code are available upon request from the Lead Author.

