## Supplemental Figures and Legends for "Sleep promotes downward firing rate homeostasis"

### Figure S1

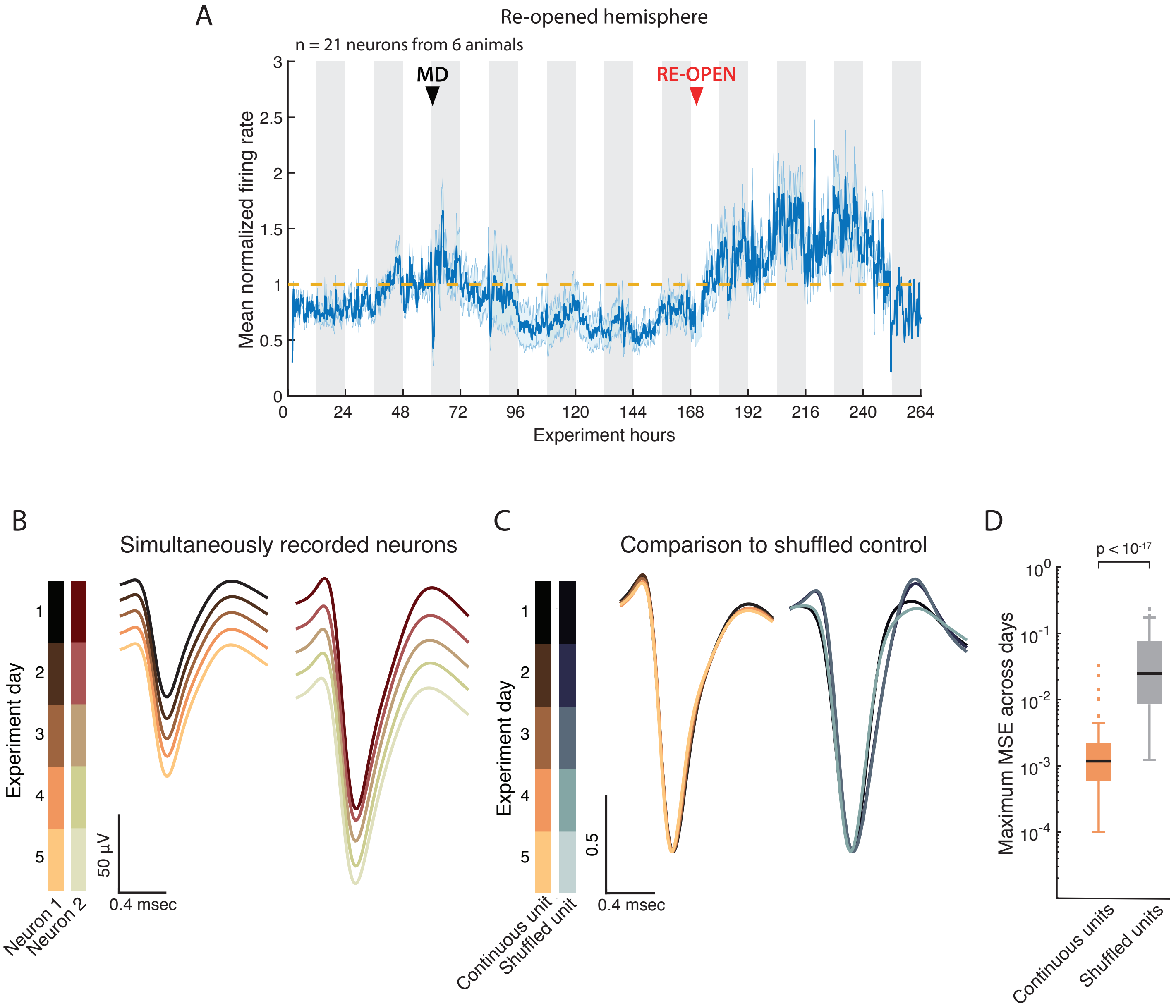

A
Re-opened hemisphere
n = 21 neurons from 6 animals
MD
RE-OPEN
Mean normalized firing rate
0
0.5
1
1.5
2
2.5
3
0
24
48
72
96
120
144
168
192
216
240
264
Experiment hours
B
Simultaneously recorded neurons
Experiment day
1
2
3
4
5
50 µV
0.4 msec
Neuron 1
Neuron 2
C
Comparison to shuffled control
Experiment day
1
2
3
4
5
0.5
0.4 msec
Continuous unit
Shuffled unit
D
p < 10^-17
Maximum MSE across days
10^0
10^-1
10^-2
10^-3
10^-4
Continuous units
Shuffled units

### Figure S2

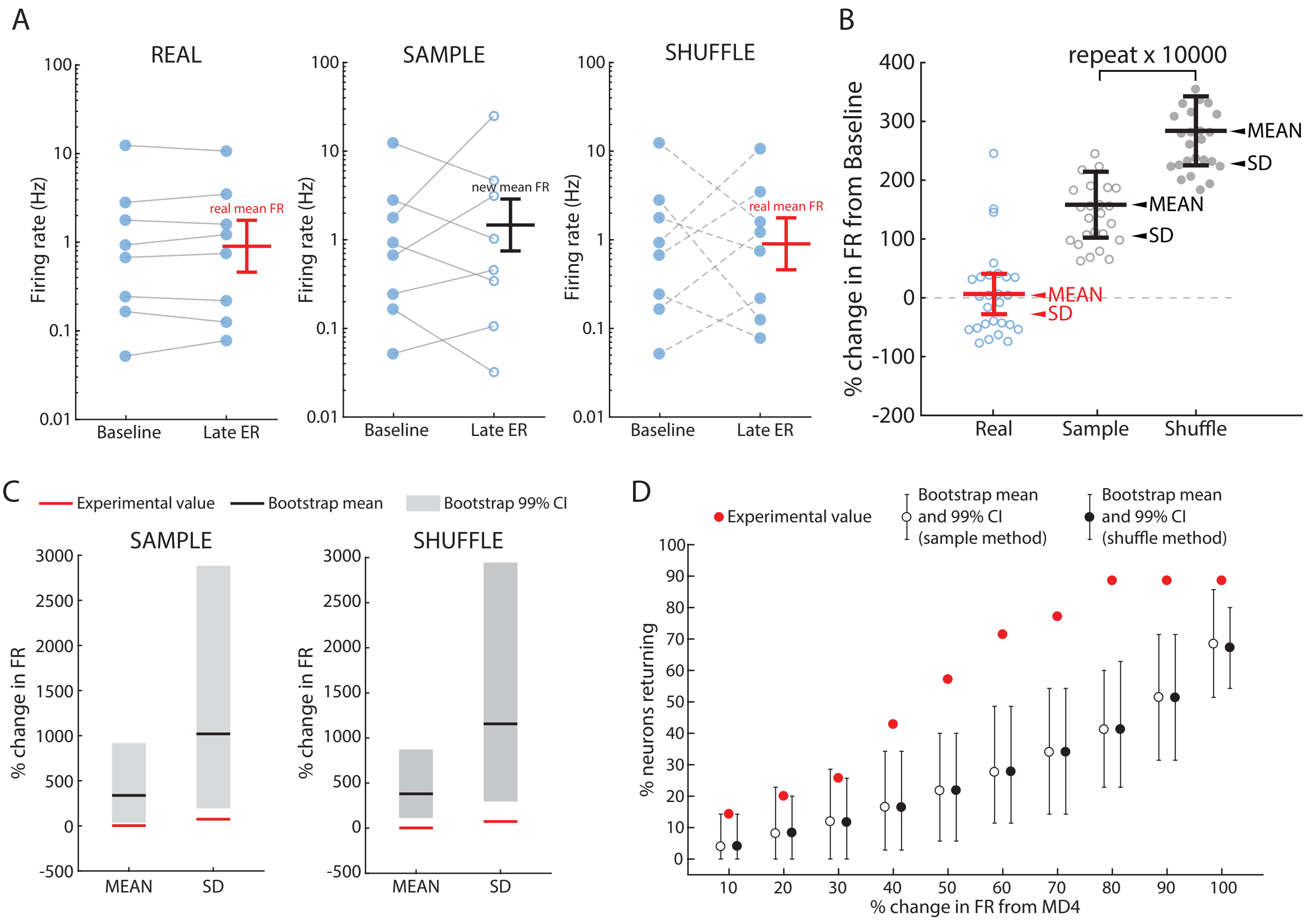
A
REAL
SAMPLE
SHUFFLE
Firing rate (Hz)
Baseline
Late ER
real mean FR
new mean FR
B
repeat x 10000
% change in FR from Baseline
MEAN
SD
Real
Sample
Shuffle
C
Experimental value
Bootstrap mean
Bootstrap 99% CI
% change in FR
D
Experimental value
Bootstrap mean and 99% CI (sample method)
Bootstrap mean and 99% CI (shuffle method)
% neurons returning
% change in FR from MD4

### Figure S3

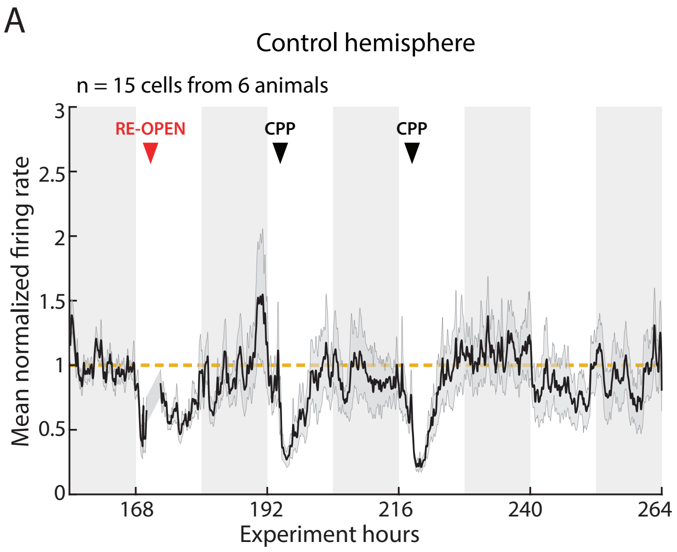
A
Control hemisphere
n = 15 cells from 6 animals
RE-OPEN
CPP
CPP
Mean normalized firing rate
3
2.5
2
1.5
1
0.5
0
168
192
216
240
264
Experiment hours

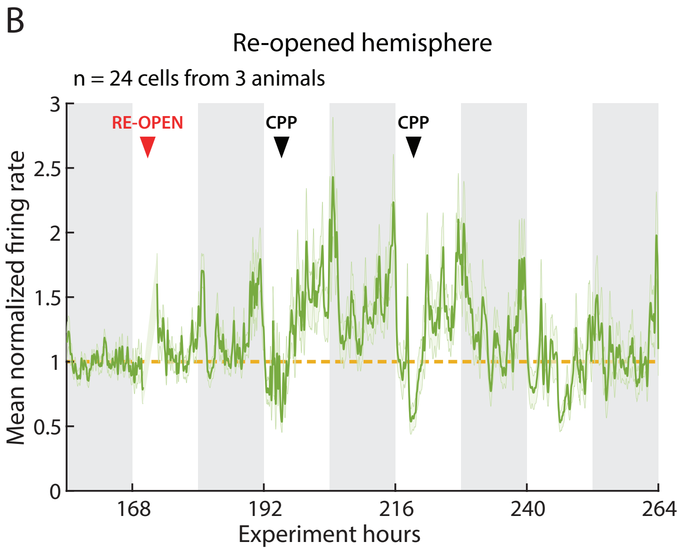
B
Re-opened hemisphere
n = 24 cells from 3 animals
RE-OPEN
CPP
CPP
Mean normalized firing rate
3
2.5
2
1.5
1
0.5
0
168
192
216
240
264
Experiment hours

### Figure S4

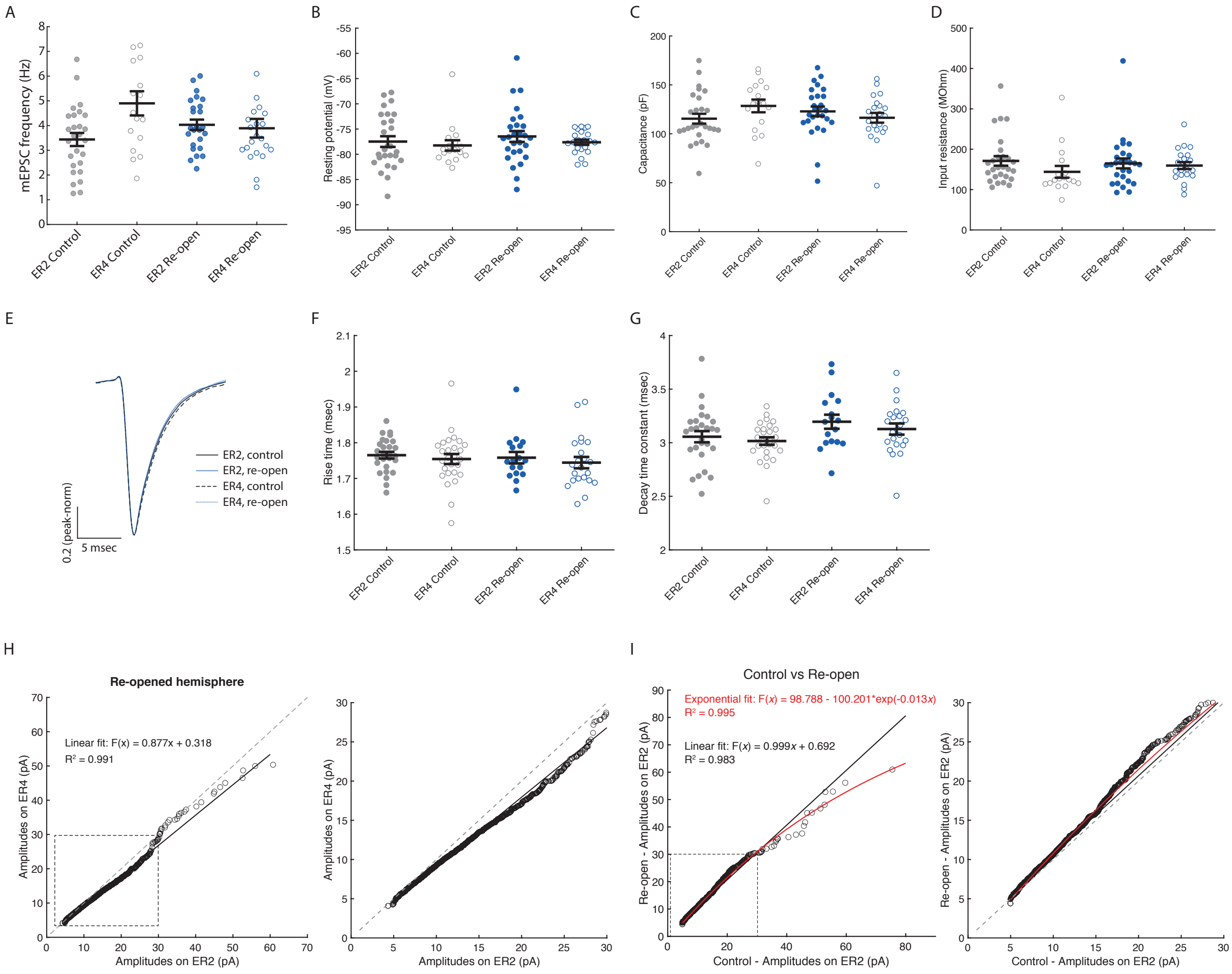
A
mEPSC frequency (Hz)
ER2 Control
ER4 Control
ER2 Re-open
ER4 Re-open
B
Resting potential (mV)
C
Capacitance (pF)
D
Input resistance (MOhm)
E
ER2, control
ER2, re-open
ER4, control
ER4, re-open
0.2 (peak-norm)
5 msec
F
Rise time (msec)
G
Decay time constant (msec)
H
Re-opened hemisphere
Linear fit: F(x) = 0.877x + 0.318
R² = 0.991
Amplitudes on ER4 (pA)
Amplitudes on ER2 (pA)
I
Control vs Re-open
Exponential fit: F(x) = 98.788 - 100.201*exp(-0.013x)
R² = 0.995
Linear fit: F(x) = 0.999x + 0.692
R² = 0.983
Re-open - Amplitudes on ER2 (pA)
Control - Amplitudes on ER2 (pA)

### Figure S5

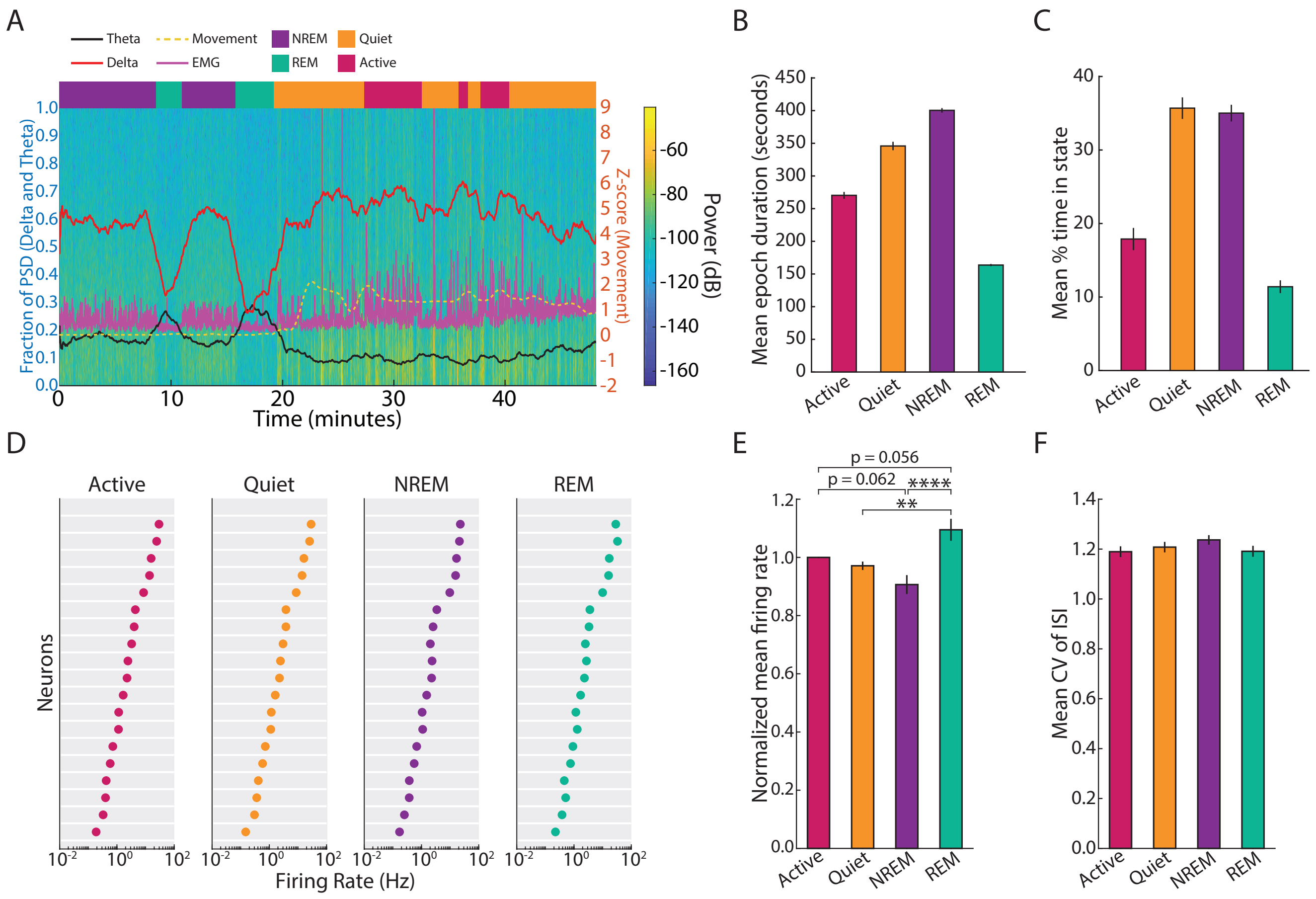
A
Theta
Delta
Movement
EMG
NREM
REM
Quiet
Active
Fraction of PSD (Delta and Theta)
Z-score (Movement)
Power (dB)
Time (minutes)
B
Mean epoch duration (seconds)
C
Mean % time in state
Active
Quiet
NREM
REM
D
Active
Quiet
NREM
REM
Neurons
Firing Rate (Hz)
E
Normalized mean firing rate
p = 0.056
p = 0.062
****
**
F
Mean CV of ISI

### Figure S6

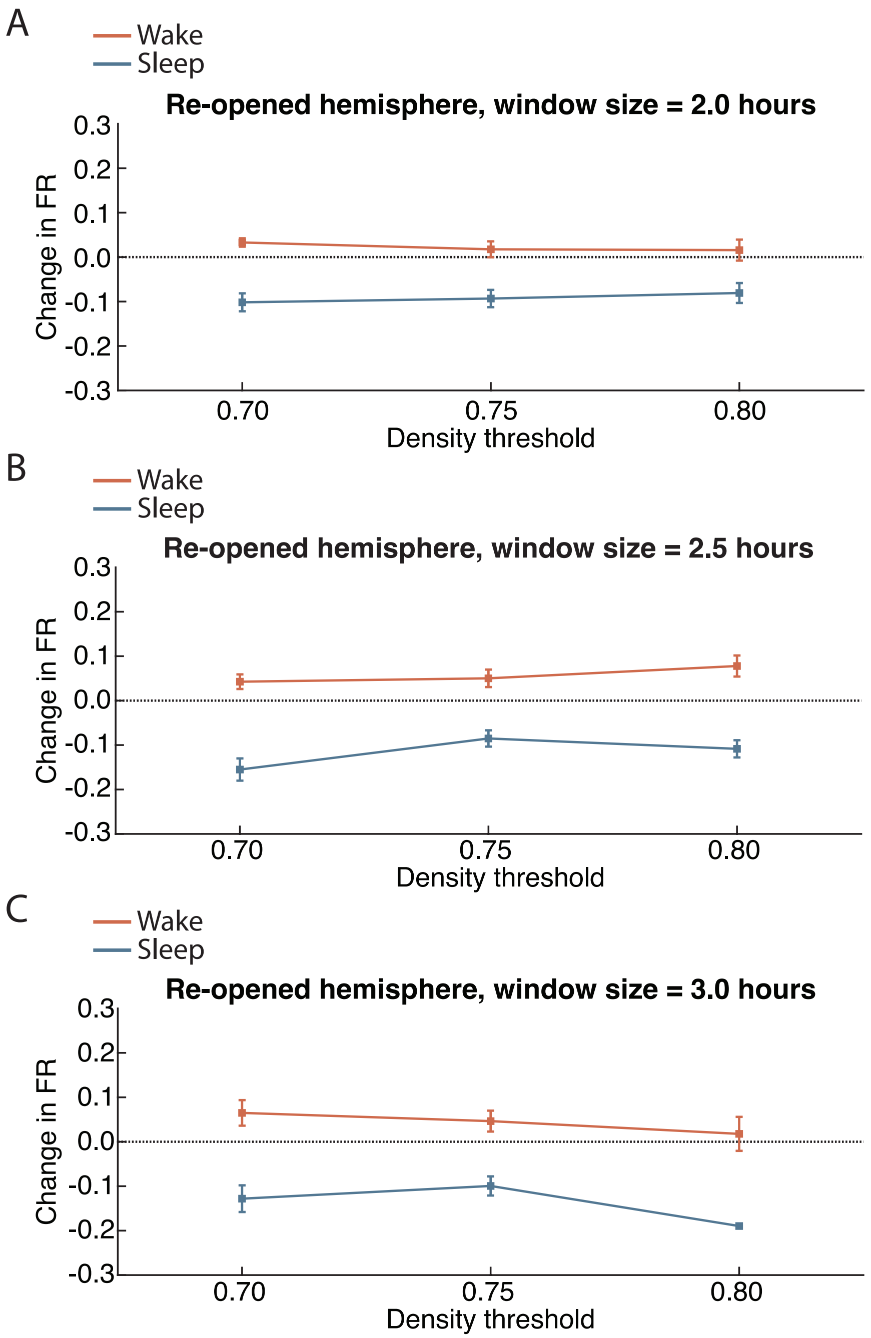
A
Wake
Sleep
Re-opened hemisphere, window size = 2.0 hours
Change in FR
0.3
0.2
0.1
0.0
-0.1
-0.2
-0.3
0.70
0.75
0.80
Density threshold
B
Wake
Sleep
Re-opened hemisphere, window size = 2.5 hours
Change in FR
0.3
0.2
0.1
0.0
-0.1
-0.2
-0.3
0.70
0.75
0.80
Density threshold
C
Wake
Sleep
Re-opened hemisphere, window size = 3.0 hours
Change in FR
0.3
0.2
0.1
0.0
-0.1
-0.2
-0.3
0.70
0.75
0.80
Density threshold

#### SUPPLEMENTAL FIGURE LEGENDS

***Figure S1: Continuous recordings of V1 neuronal activity in freely behaving rats.*** **A.** Average FR, normalized to baseline (hours 36-60), for all neurons recorded continuously in the deprived/re-opened hemisphere for 11 days throughout baseline, MD and ER. Yellow dashed line indicates mean baseline FR. Labeled arrows indicate time of MD and ER. White/gray boxes in background denote 12-hour periods of light/dark. **B.** Daily average waveforms (WFs) for two RSUs (putative excitatory neurons) recorded simultaneously on the same wire for 5 days (MD4 to ER4). Note similarity of WFs across days, and discriminability between the two neurons. **C.** Example peak-scaled daily average WFs for a continuously recorded neuron (left) and a shuffled random unit (right). See STAR methods for details ("Automated spike extraction, clustering and sorting"). **D.** Result of bootstrap analysis (see STAR methods for details). Maximum mean-squared-error (MSE) between daily peak-scaled average WFs for continuous vs shuffled random units. Two-sample t-test, $p < 10^{-17}$.

***Figure S2: neurons recover to their initial baseline FR after ER-induced overshoot.*** **A.** Diagram illustrating the two bootstrap methods used (see STAR methods, "Bootstrap analysis of FR recovery", for details). Reference data (left) are the baseline and late ER distribution of FRs in the re-opened hemisphere (data points shown here are simulated, for illustration purposes only, see Figure 1 for real data). Red line and errorbars denote real mean FR of the distribution at late ER. The "sample" bootstrap strategy (middle) consists in sampling at random from the empirical distribution of FRs at late ER, and randomly matching to baseline values. Note that this results in a mean population FR different than that of real data. The "shuffle" bootstrap strategy (right) does not resample the late ER distribution, but simply shuffles the matching between baseline and late ER data points. This preserves the real mean population FR, but not the % change of individual neurons. **B.** Quantification of bootstrap analysis. For each sampling or shuffling iteration the % change in FR from baseline is computed for each neuron, and the resulting mean and standard deviation (SD) of the distribution are calculated. This is repeated over 10,000 iterations to obtain confidence intervals (CI) for the mean and SD. **C.** Bootstrap analysis results for sample (left) and shuffle (right) methods. The red line indicates real mean and SD for the data in Figure 1F. Black lines indicate average values over 10,000 bootstrap iterations. Gray shaded area indicates the 99% CI for mean and SD. **D.** Estimate of the FR range over which neurons vary their FR, compared to bootstrap control (see STAR methods, "Bootstrap analysis of FR recovery", for details). Red dots indicate experimental value, and means and 99% CIs for sample (open circles) and shuffle (filled circles) methods are shown.

***Figure S3: Acute effect of CPP injections on FRs in V1.*** **A.** Average baseline-normalized FR for neurons recorded in the control hemisphere of animals injected with CPP during recovery of FR after ER. CPP injections (labeled black arrows) briefly depress FRs, but cause no long-lasting effects in the control hemisphere. Dashed yellow line indicates baseline FR. **B.** As in A, but for neurons recorded in the re-opened hemisphere. CPP injections (labeled black arrows) have the same acute effect, but FRs still show overshoot and recovery following ER.

***Figure S4:*** **ER does not cause change in passive properties or mEPSC kinetics. A – D.** Passive neuronal properties for all cells recorded in each condition. No change in mEPSC frequency (A), resting membrane potential (B), cell capacitance (C) or input resistance (D). Kruskal-Wallis test with Tukey-Kramer post-hoc, no significant results. **E.** Peak-scaled average mEPSC waveforms for each condition, overlaid. **F, G.** Summary of waveform kinetics for all cells in each condition. No change in rise time (F) or decay time constant (G). **H.** Sorted mEPSC amplitudes at ER2 vs ER4 in the re-opened hemisphere, showing scaling relationship. Dashed line is the unity line, solid line indicates linear fit and each open circle is one mEPSC event. Right, zoomed-in view of the plot region inside the black dashed rectangle in the left plot. **I.** Sorted mEPSC amplitudes in the control vs re-opened hemispheres at ER2. In this case the data are better fit by an exponential than a linear function. Scaling the control distribution by a linear function does not recover the re-opened distribution (see Figure 3F). Black solid line indicates linear fit, red solid line indicates exponential fit and open circles are individual mEPSC events. Right, zoomed-in view of the plot region inside the black dashed rectangle in the left plot.

***Figure S5: Behavioral state scoring and FR differences across states.*** **A.** Example behavioral scoring for a ~30-minute period. Background spectrogram is the LFP power between 0.3 and 15 Hz (colorbar on the right). Red and black line represent the fraction of power in the delta (0.3 – 4 Hz) and theta (5 – 8 Hz) bands (plotted on left y-axis). The yellow dashed lines represent movement values, z-scored to the 60-minute block being scored (plotted on right y-axis). Solid violet line represents the EMG data (plotting axes not shown). Colored rectangles at the top show the scored state after manual correction of random forest classifier output (NREM sleep, REM sleep, Active wake or Quiet wake). **B.** Mean epoch duration for each behavioral state for all animals. **C.** Average percent time in each state for all animals. **D.** Mean FR of every RSU recorded in the control hemisphere in each state, ranked by mean FR in active wake. **E.** Average FR in each state for RSUs recorded in the control hemisphere, normalized for each neuron to the mean FR in active wake. One-way ANOVA with Tukey-Kramer post-hoc, ** $p = 0.007$, **** $p < 10^{-4}$. **F.** Mean coefficient of variation (CV) of the inter-spike interval (ISI) in each state.

***Figure S6: Sleep- and wake-dense analysis result does not depend on chosen parameters.*** **A – C.** Mean change in FR for RSUs in the re-opened hemisphere in sleep- and wake-dense windows for different density thresholds. The analysis was repeated for 2-hour (A), 2.5-hour (B) and 3-hour (C) windows. Density threshold is minimum % time in sleep- or wake- for a window to be considered sleep- or wake-dense. Results are presented as mean ± SEM.
